# The Seroprevalence of SARS-CoV-2 in Europe: A Systematic Review

**DOI:** 10.1101/2021.04.12.439425

**Authors:** Natasha Marcella Vaselli, Daniel Hungerford, Ben Shenton, Arwa Khashkhusha, Nigel A. Cunliffe, Neil French

## Abstract

**Background:** A year following the onset of the COVID-19 pandemic, new infections and deaths continue to increase in Europe. Serological studies, through providing evidence of past infection, can aid understanding of the population dynamics of SARS-CoV-2 infection.

**Objectives:** This systematic review of SARS-CoV-2 seroprevalence studies in Europe was undertaken to inform public health strategies including vaccination, that aim to accelerate population immunity.

**Methods:** We searched the databases Web of Science, MEDLINE, EMBASE, SCOPUS, Cochrane Database of Systematic Reviews and grey literature sources for studies reporting seroprevalence of SARS-CoV-2 antibodies in Europe published between 01/12/2019 - 30/09/20. We provide a narrative synthesis of included studies. Studies were categorized into subgroups including healthcare workers (HCWs), community, outbreaks, pregnancy and children/school. Due to heterogeneity in other subgroups, we only performed a random effects meta-analysis of the seroprevalence amongst HCWs stratified by their country.

**Results:** 109 studies were included spanning 17 European countries, that estimated the seroprevalence of SAR-CoV2 from samples obtained between November 2019 – August 2020. A total of 53/109 studies included HCWs with a reported seroprevalence among HCWs ranging from 0.7% to 45.3%, which did not differ significantly by country. In community studies significant heterogeneity was reported in the seroprevalence among different age groups and the majority of studies reported there was no significant difference by gender.

**Conclusion:** This review demonstrates a wide heterogeneity in reported seroprevalence of SARS-CoV-2 antibodies between populations. Continued evaluation of seroprevalence is required to understand the impact of public health measures and inform interventions including vaccination programmes.

## Introduction

On 11^th^ March 2020 the World Health Organization (WHO) declared the spread of novel SARS-CoV-2 as a pandemic (1). SARS-CoV2 is thought to spread mainly by respiratory droplets, while some evidence also suggests spread via fomites and aerosols (2–4). SARS-CoV2 causes varying degrees of illness from mild symptoms including fatigue and myalgia to acute respiratory failure and death (5). As the pandemic unfolded evidence emerged that a large number of individuals are asymptomatic with SARS-CoV2 infection (6, 7).

In order to control the spread of SARS-CoV-2 it is important to understand the extent to which different populations have already been exposed to the virus, especially as a large number of infections are asymptomatic. Many countries, organizational bodies and facilities have turned to mass testing to estimate the spread of infection and inform public health measures (8, 9). One such testing strategy is by nasal and throat swabbing to detect viral RNA which has recently been piloted in England (10). A further method is mass testing of the population for antibodies against SARS-CoV-2 (11). Several tests for IgG, IgA and IgM antibodies against SARS-CoV2 have recently been developed. These broadly include enzyme linked immunosorbent assays, chemiluminescence immunoassays (CLIA) and point of care lateral flow assays (12).

Seroprevalence studies have been used in the past to help with outbreak responses. During a recent Ebola outbreak, seroprevalence studies were performed to gain further information on the immune response and immune protection (13). Seroprevalence studies have also been used for infections such rubella, mumps and measles to map resurgence and to gain further information on how public health strategies can target high risk populations (14). Moreover, seroprevalence studies provide valuable information on vaccination strategies to achieve herd immunity.

By estimating the seroprevalence in different populations we can use the results to understand transmission dynamics, herd immunity and the immune response over time. These studies can help to guide the public health response to ultimately help prevent the spread of SARS-CoV2. Here we present the results of a systematic review on the seroprevalence of SARS-CoV2 in Europe.

## Methods

### Search strategy and selection criteria

This systematic review and meta-analysis adhere to the Preferred Reporting Items for Systematic Reviews and Meta-Analyses (PRISMA) guidelines. The protocol was registered on the University of York database for Prospectively Registered Systematic Reviews (PROSPERO: 2020 CRD42020212149). We systematically searched electronic data sources from (01/12/2019) until (30/09/20) using search terms on seroprevalence and SARS-CoV-2. We searched the following electronic databases: Web of Science, MEDLINE, EMBASE, SCOPUS and Cochrane Database of Systematic Reviews. We also searched database search engine EBSCO to search the following databases EBSCOhost e-book collection, biomedical reference collection, CINAHL plus and MEDLINE Complete. We conducted a secondary search by searching the reference lists of included studies for relevant articles.

Furthermore, we searched the grey literature. Firstly, we searched pre-print articles in the electronic database search engine EPMC to search pre-print databases including MedRXIV and BioRXIV. Secondly, we then used the database OpenGrey to search research reports, doctoral dissertations, conference papers and other forms of grey literature. Thirdly we searched the websites of national and international health agencies for reports relating to the seroprevalence of SARS-CoV-2 (World Health Organization, European Centres for Disease Control, Public Health England, Department of Health and Social Care in UK). Finally, we conducted a google search for further government reports.

Search terms were developed alongside a health science librarian.

| Search Strategy |  |
| --- | --- |
| 1 | COVID-19 OR SARS CoV-2 |
| 2 | antibod* OR immun* |
| 3 | Seroprevalence |
| 4 | 1 AND 2 AND 3 |
| 5 | animals NOT humans |
| 6 | 4 NOT 5 |

Studies were included if they were written in English and published between 1/12/19-30/09/20 and were cross-sectional or cohort studies. Vaccine evaluations and randomised controlled trials were excluded.

### Data extraction

Titles and abstracts of retrieved studies were de-duplicated and screened independently by two reviewers to identify if they met the inclusion criteria. Screened references then underwent full text review by two reviewers independently. Any disagreement between them over the eligibility of studies was resolved through discussion with third reviewer.

A standardised data extraction form was used. Data were extracted on the characteristics of the study (country, date, setting, selection method, funder), antibody assay employed, specificity and sensitivity of the assay, sample characteristics (age, gender, ethnicity, co-morbidities) and prevalence. Data extraction was carried out autonomously by the reviewers and consensus was sought between the team.

### Assessment of the methodological quality of included studies

We assessed the risk of bias using an adapted version of the Hoy et al modified Risk of Bias Tool, as used by Nguyen et al (15). This is a tool designed to assess the risk of bias in population-based prevalence studies. It uses a scoring system to assess the external and internal validity of the study. Studies that score 0-3 points are classified as low risk, 4-6 high risk and 7-9 high risk. Two reviewers independently applied the criteria. Disagreements were resolved through discussion.

### Data Analysis

We provided a narrative synthesis of the findings of all included studies, study population characteristics, antibody assays used and seroprevalence estimates for each study. Studies were categorized into subgroups including those that examined health care workers (HCWs), community studies, outbreaks, seroprevalence in children/ schools and seroprevalence studies performed in pregnant women. We used Metaprop in STATA version 14, statistical software (Stata Corp. College Station, TX, USA), package to perform a random– effects meta-analysis of seroprevalence amongst health care workers (HCWs) stratified by country. We also performed a random – effects meta-analysis of seroprevalence amongst health care workers (HCWs) stratified by their risk of exposure to SARS-CoV-2 infected patients. HCWs were categorised as high risk if they worked with patients with known SARS-CoV-2 infection, medium risk if they had patient contact but without known SARS-CoV-2 infection and low risk if they had no patient contact (e.g., laboratory staff and administrative staff). If studies included participants from a mixture of risk groups they were categorised as medium risk. Heterogeneity was measured using the *I*^2^ statistic which describes the percentage of total variation due to inter-study heterogeneity. Tests of heterogeneity were undertaken within the sub-groups and for the overall meta-analysis. Sensitivity analysis was done by removing those studies with a moderate risk of bias score. This had no effect of the *I*^2^ value, so these studies were included in the final meta-analysis.

After reviewing the relevant literature, we did not perform traditional asymmetry tests and funnel plots for assessing publication bias, as the meta-analysis we conducted was a summary of proportions and not a comparison of treatments/interventions (16, 17).

## Results

The literature search yielded 1648 articles. After removing duplicates and excluding studies based on their abstract or through full text examination 109 studies were identified as eligible (Figure 1). The 109 articles spanned 17 countries in Europe and estimated the seroprevalence of SAR-CoV2 from serum samples dated from November 2019 – August 2020. The 109 articles reviewed the seroprevalence in approximately 500,000 samples; 59.7% of samples belonged to females and had an overall age range of 0 – 90+ years. Data were recorded on the type of funding the studies received; this included government funding, research grants and some studies received no external funding (table 1).

**Figure 1:**
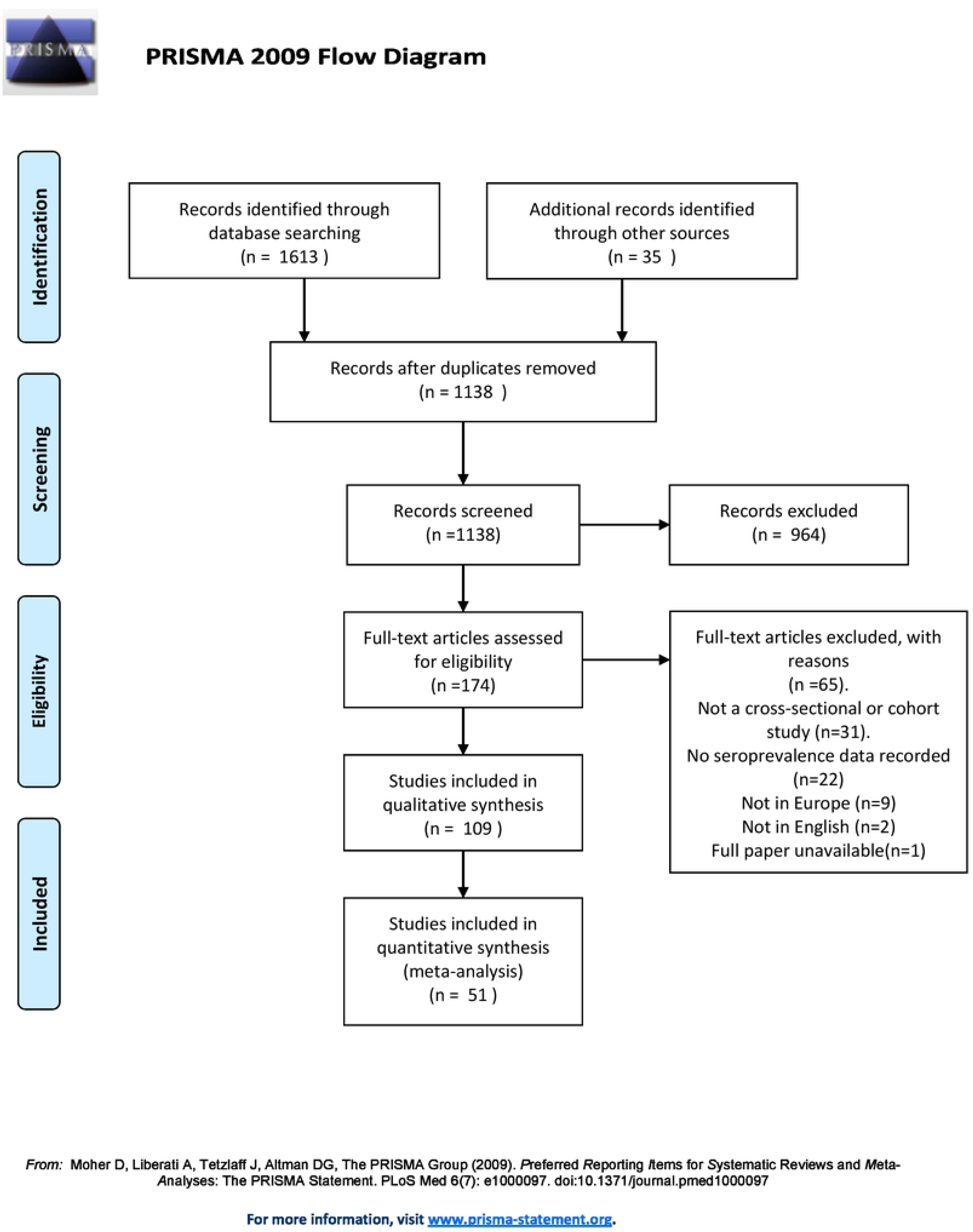
PRIMSA Flow chart

**Table 1:**
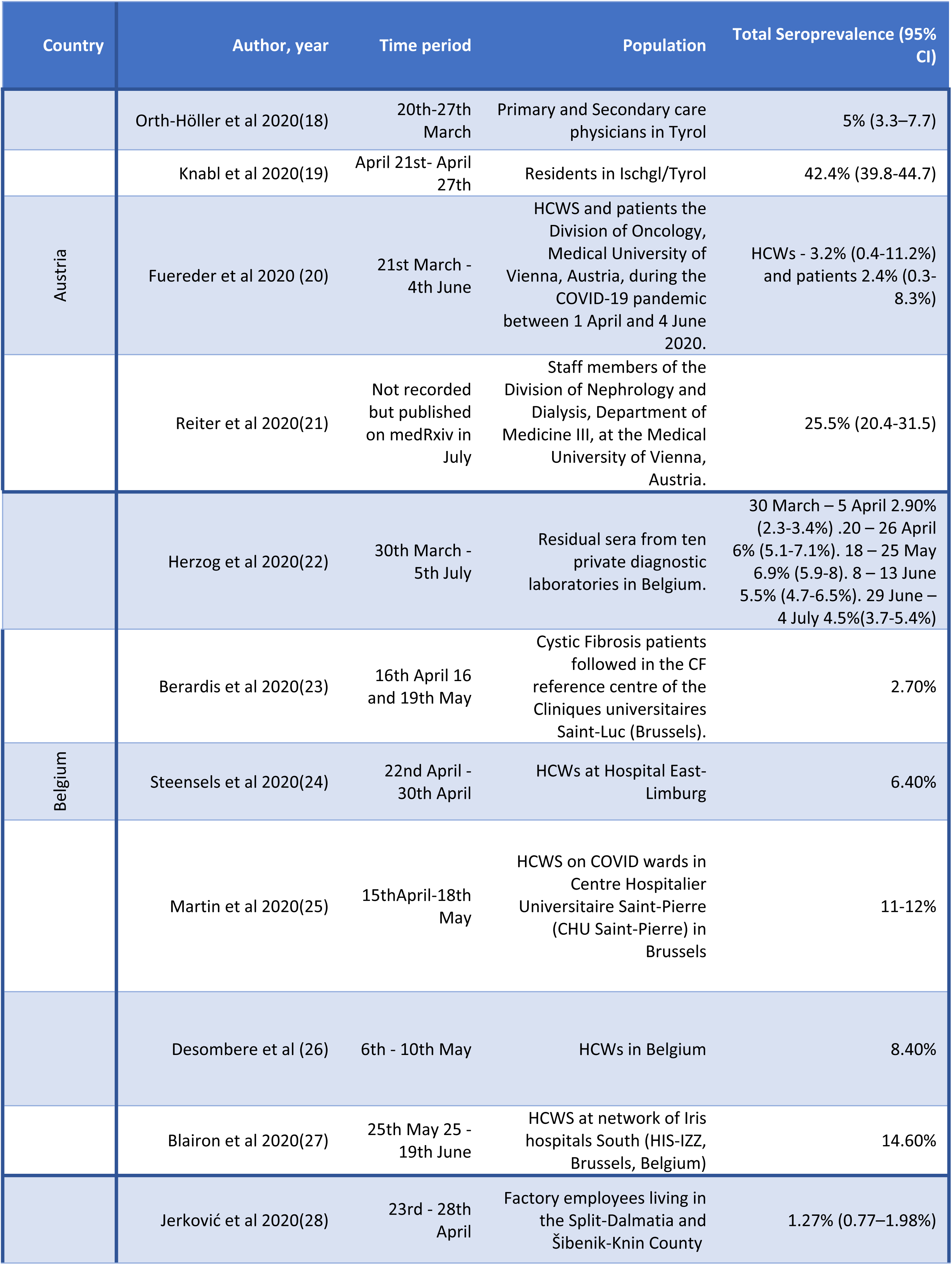
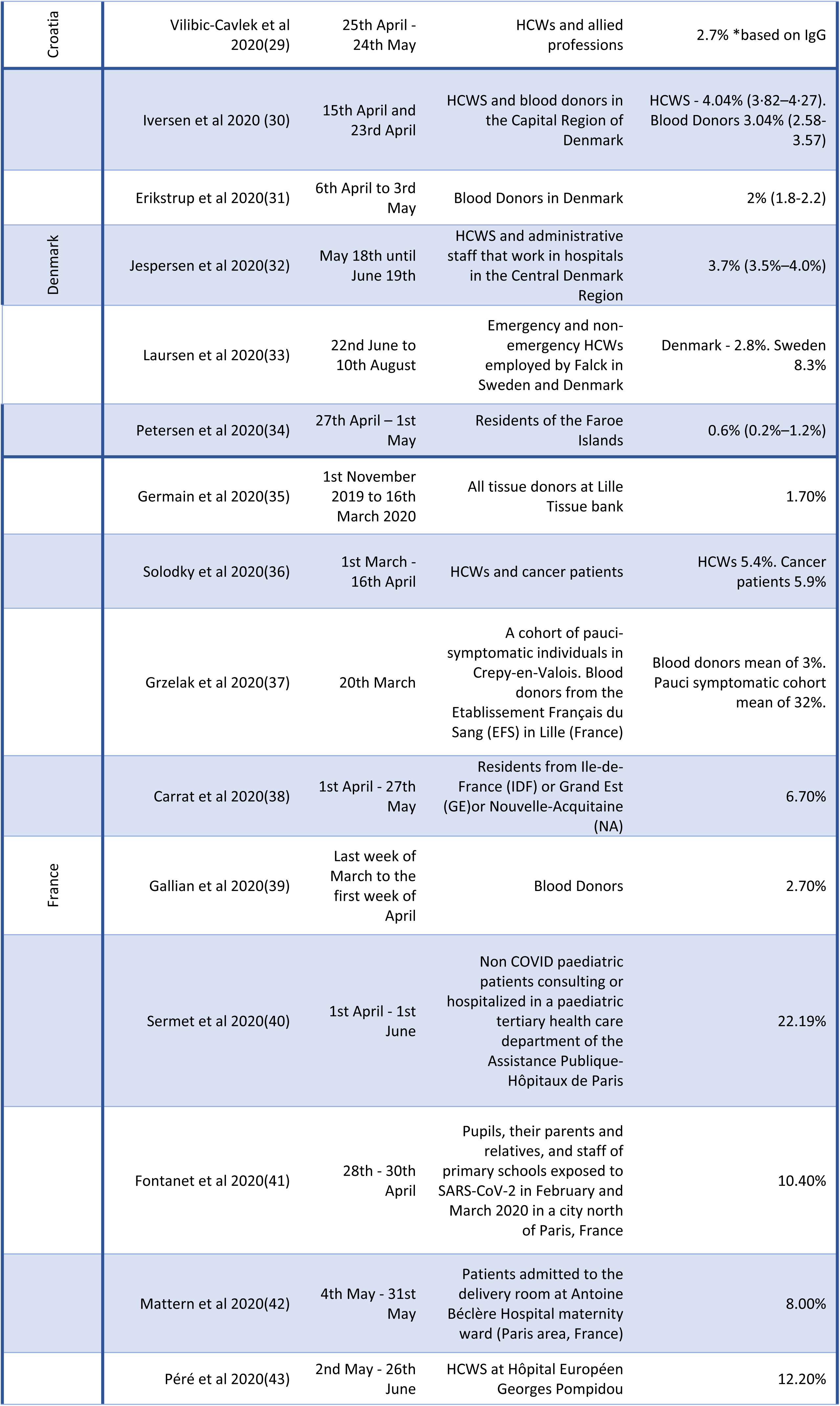
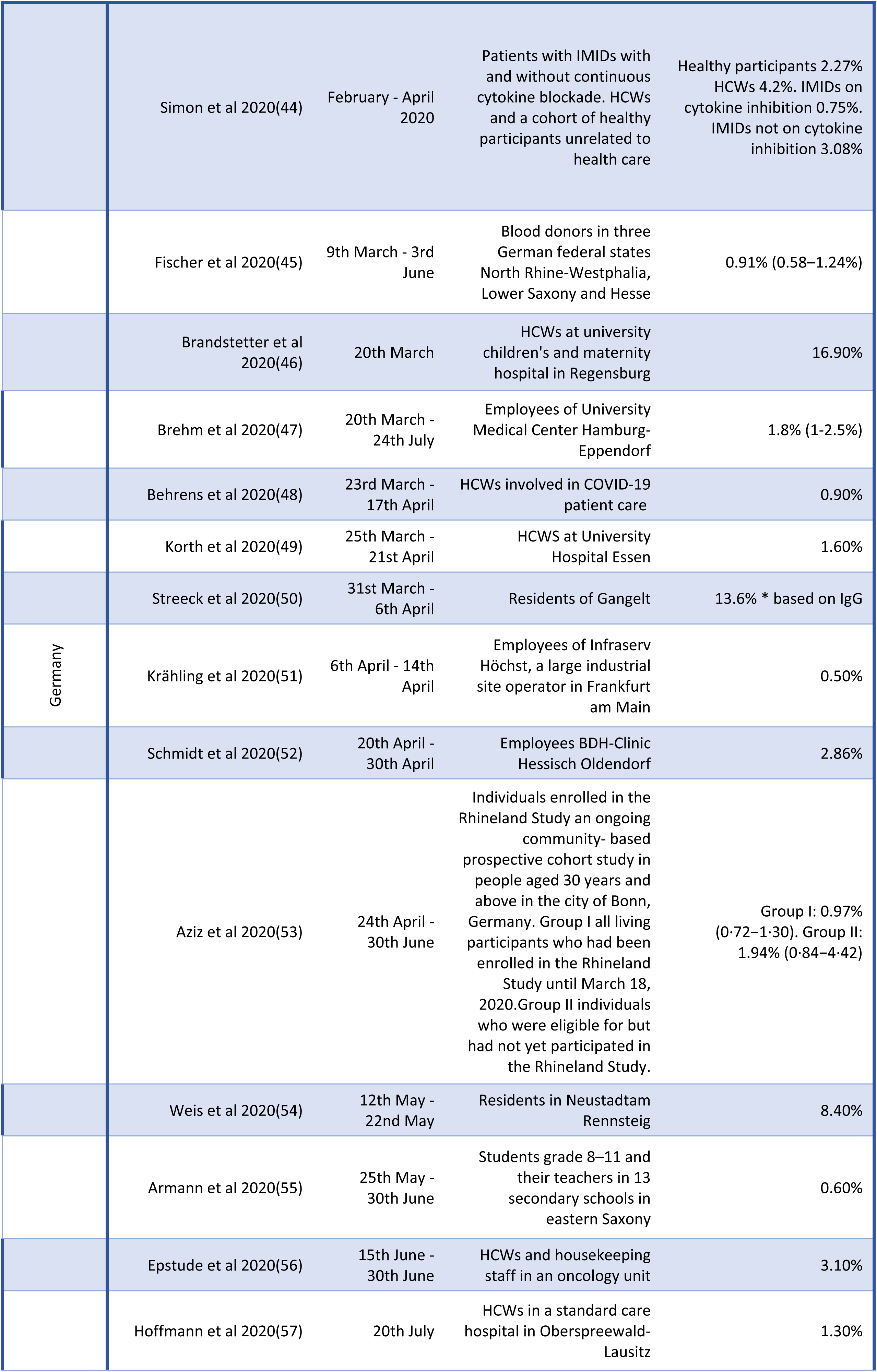
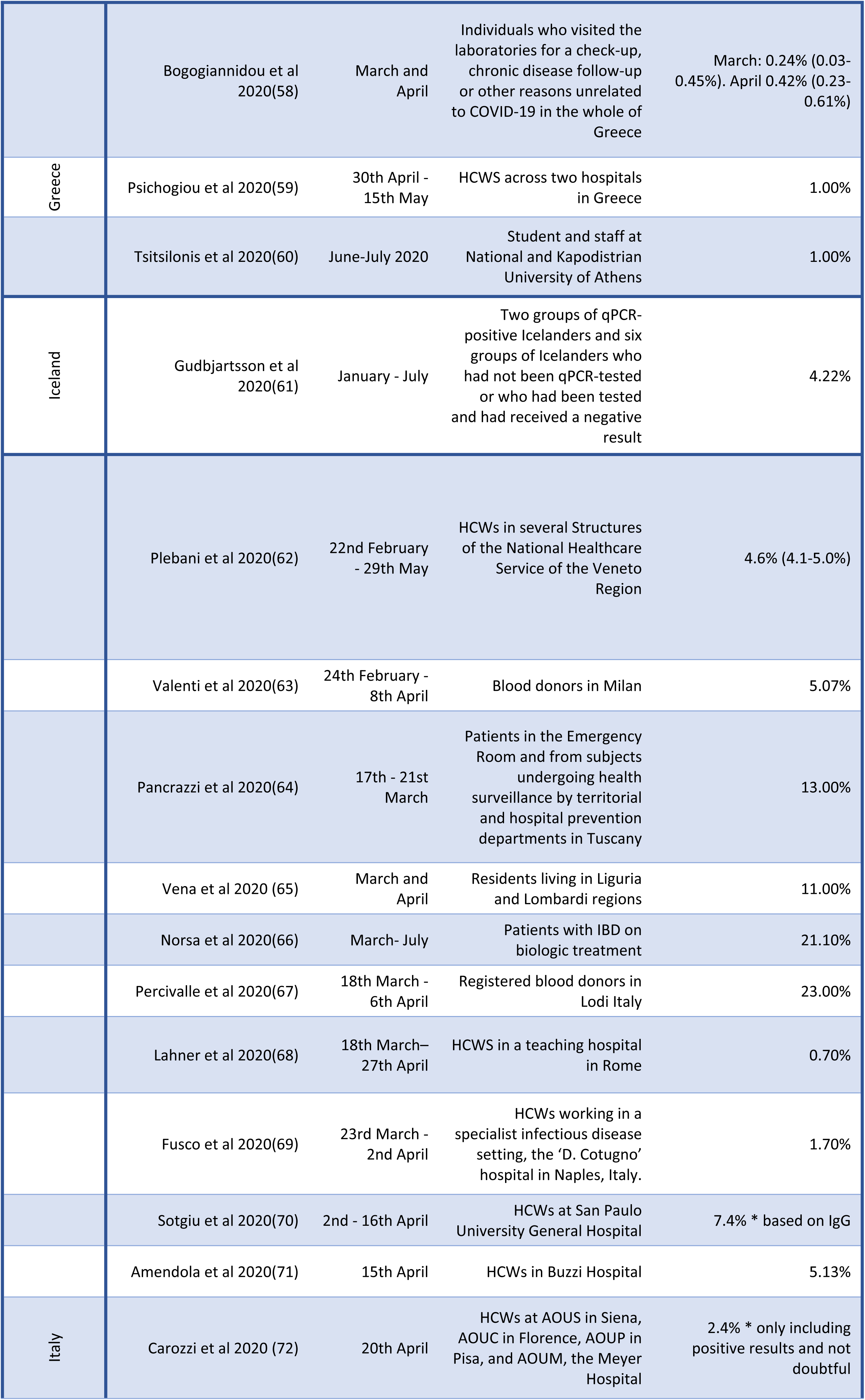
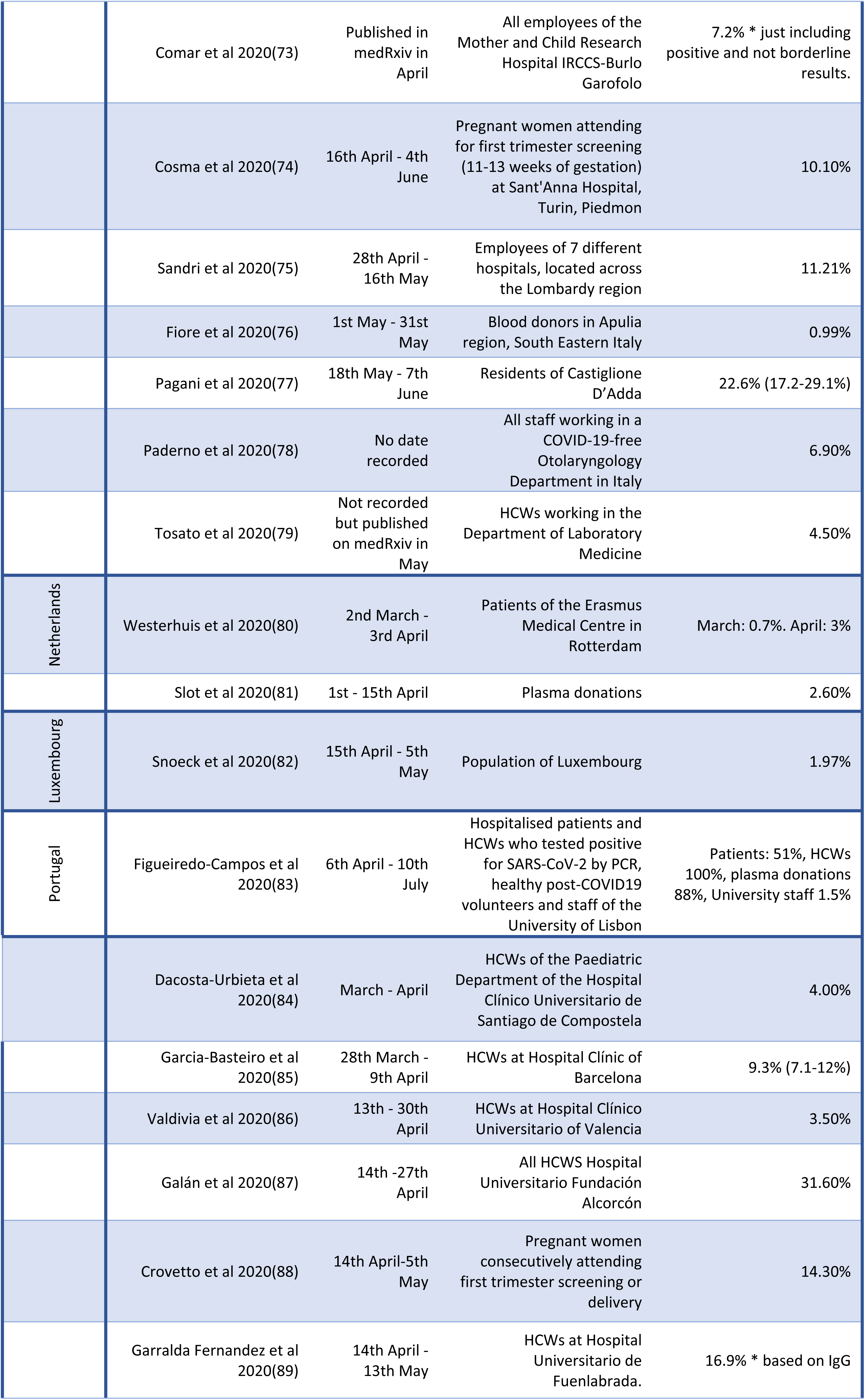
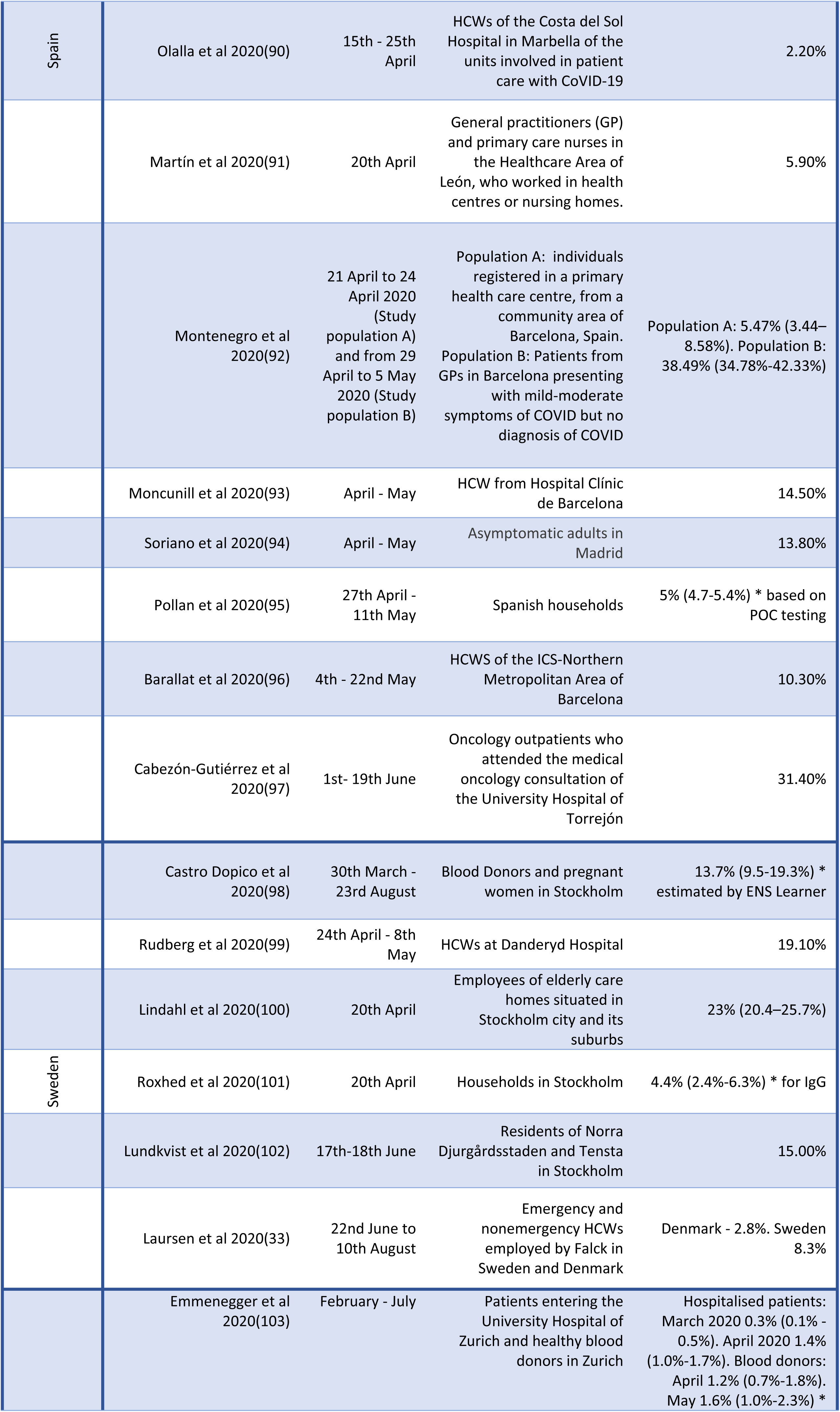
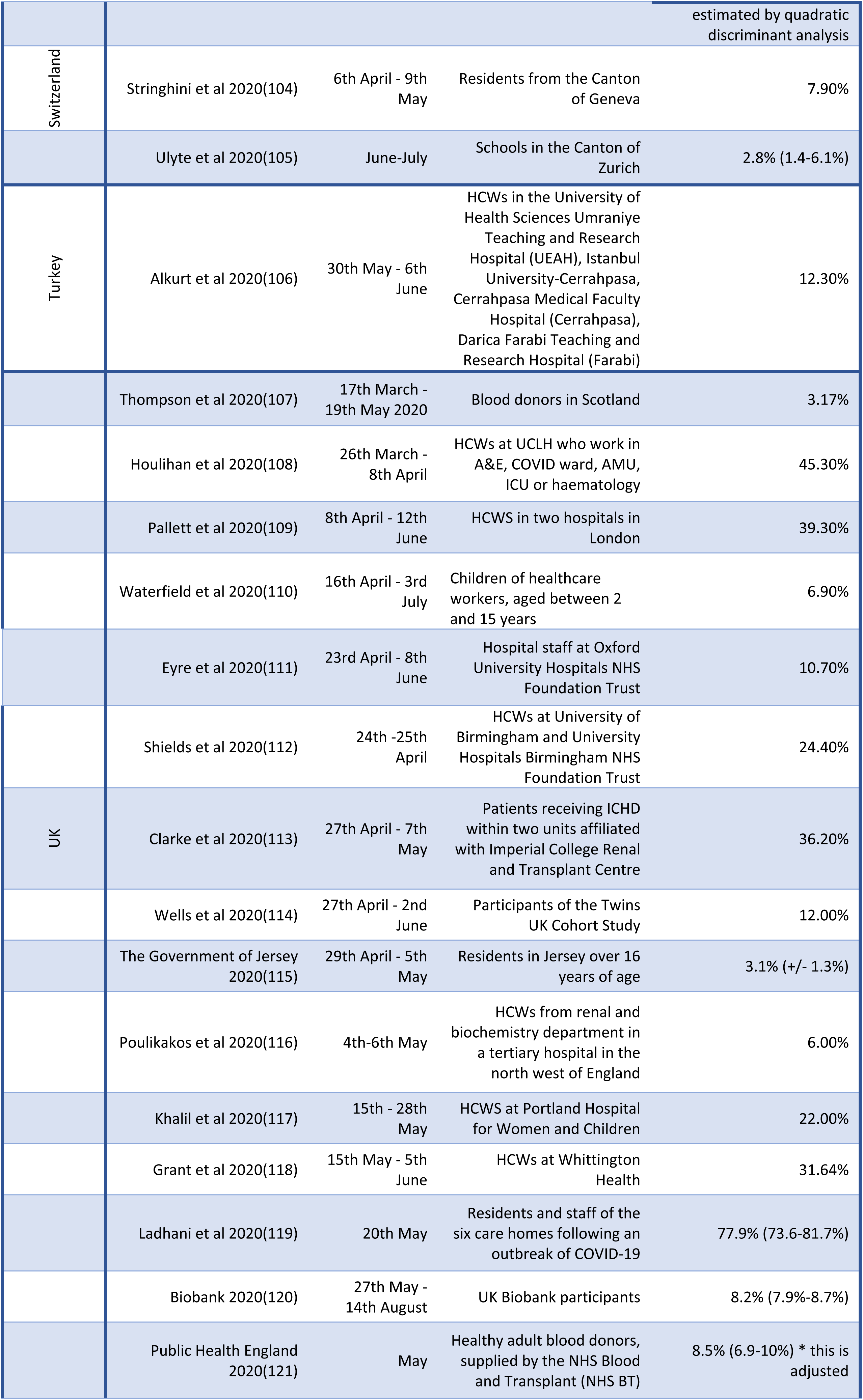
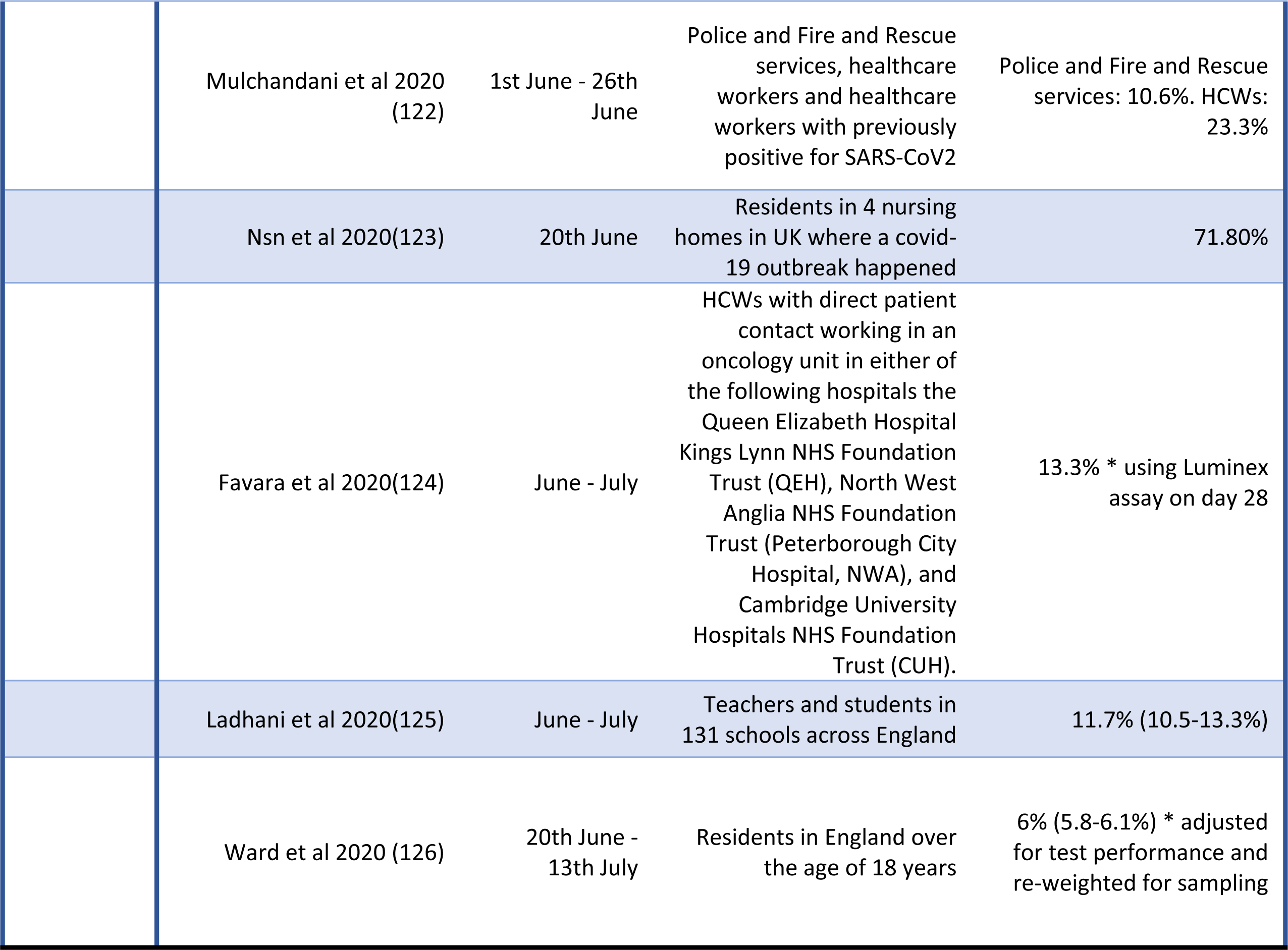
Included studies, dates of sampling, population studied and overall seroprevalence

### Assessment of the methodological quality of included studies

Each study underwent a risk of bias assessment using the modified Hoy et al risk of bias tool (15). Twenty eight of the 109 studies were deemed to be at moderate risk of bias (26,29,36,40,46,48,49,58,60,64,65,79,86,88,91,97,98,101,105,107,110,111,114,116,117,121,123,125). This was often due to lack of information about the sampling frame, selection process and non-response bias. No studies scored high risk (Table 2).

**Table 2:**
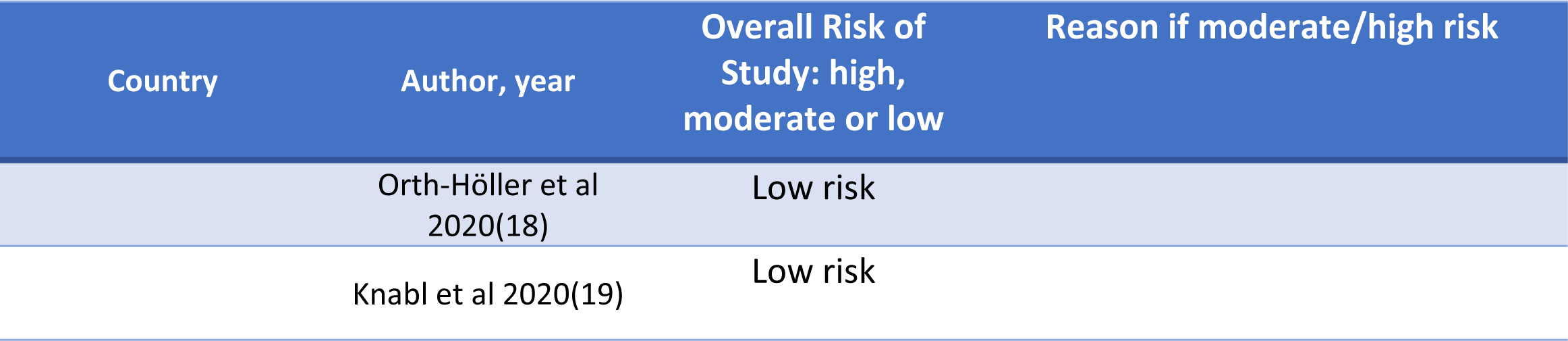
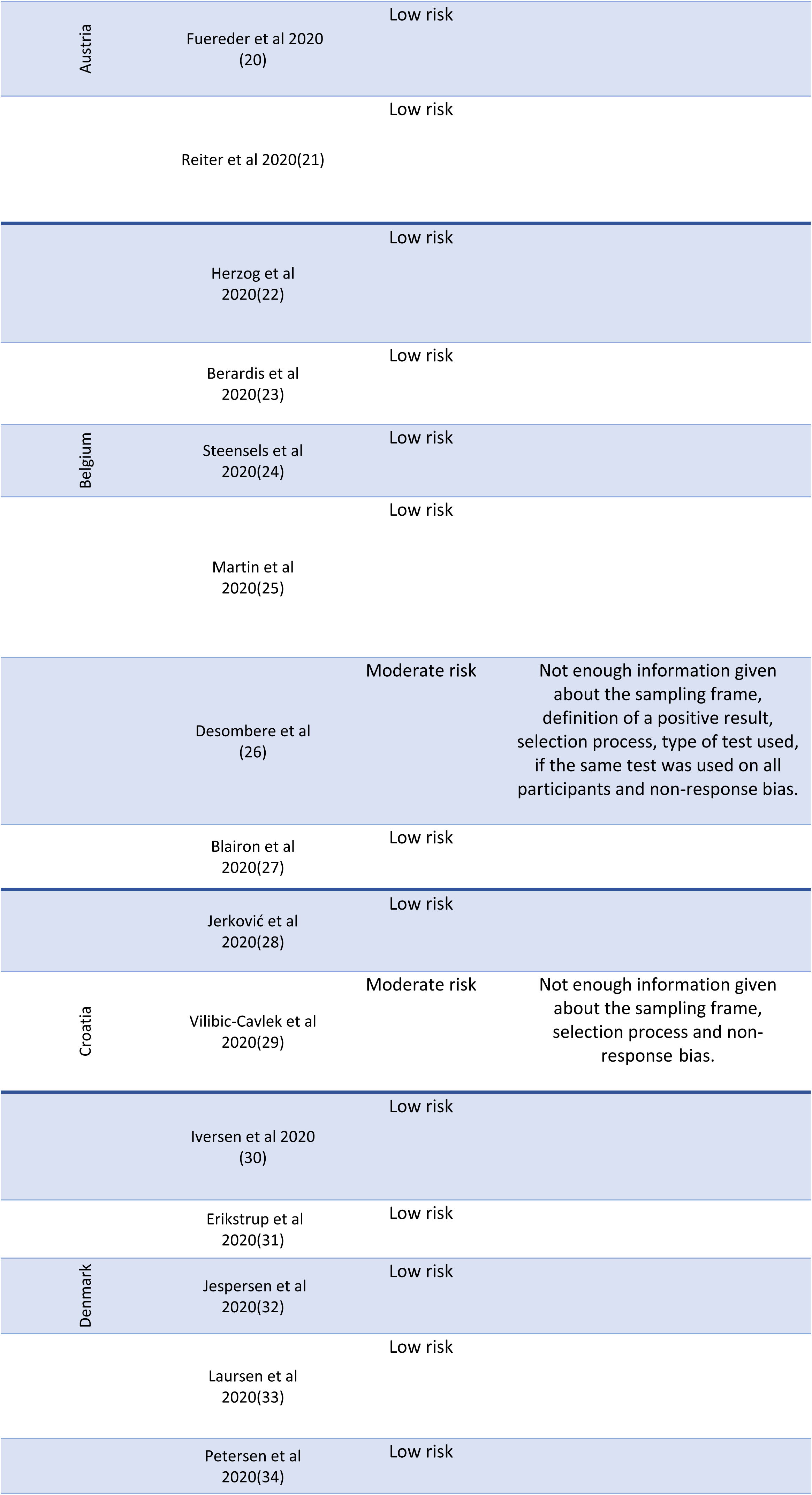
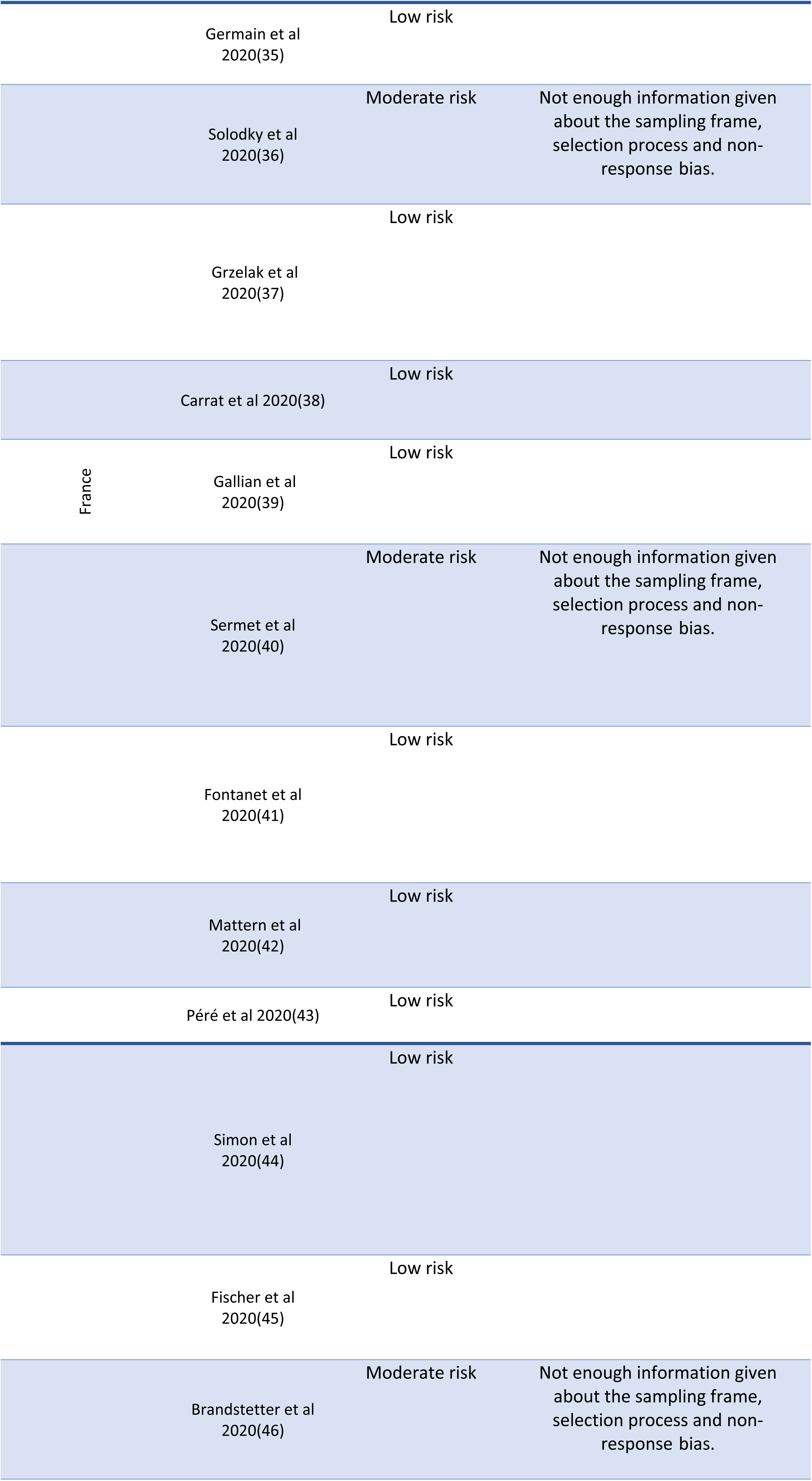
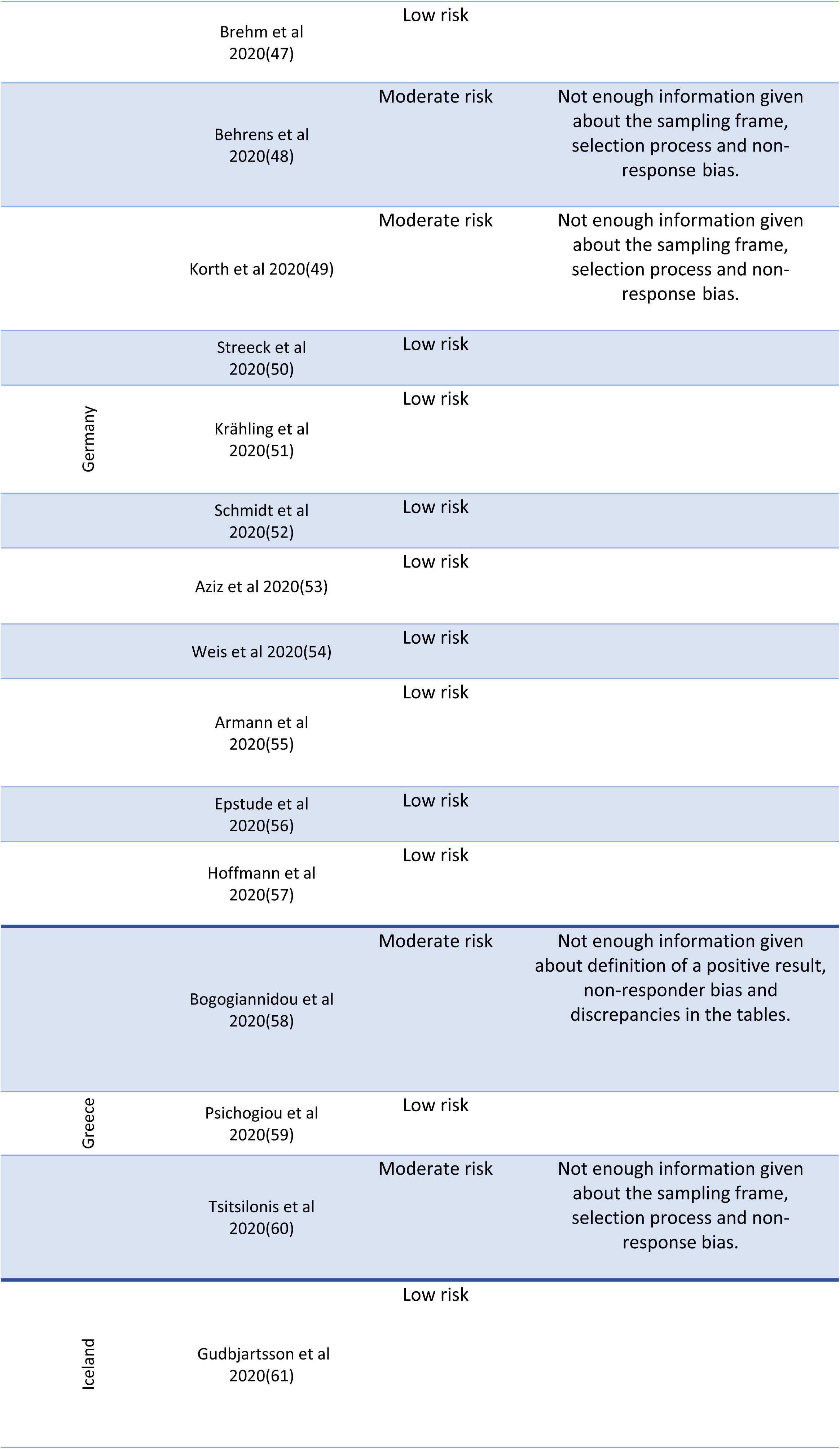
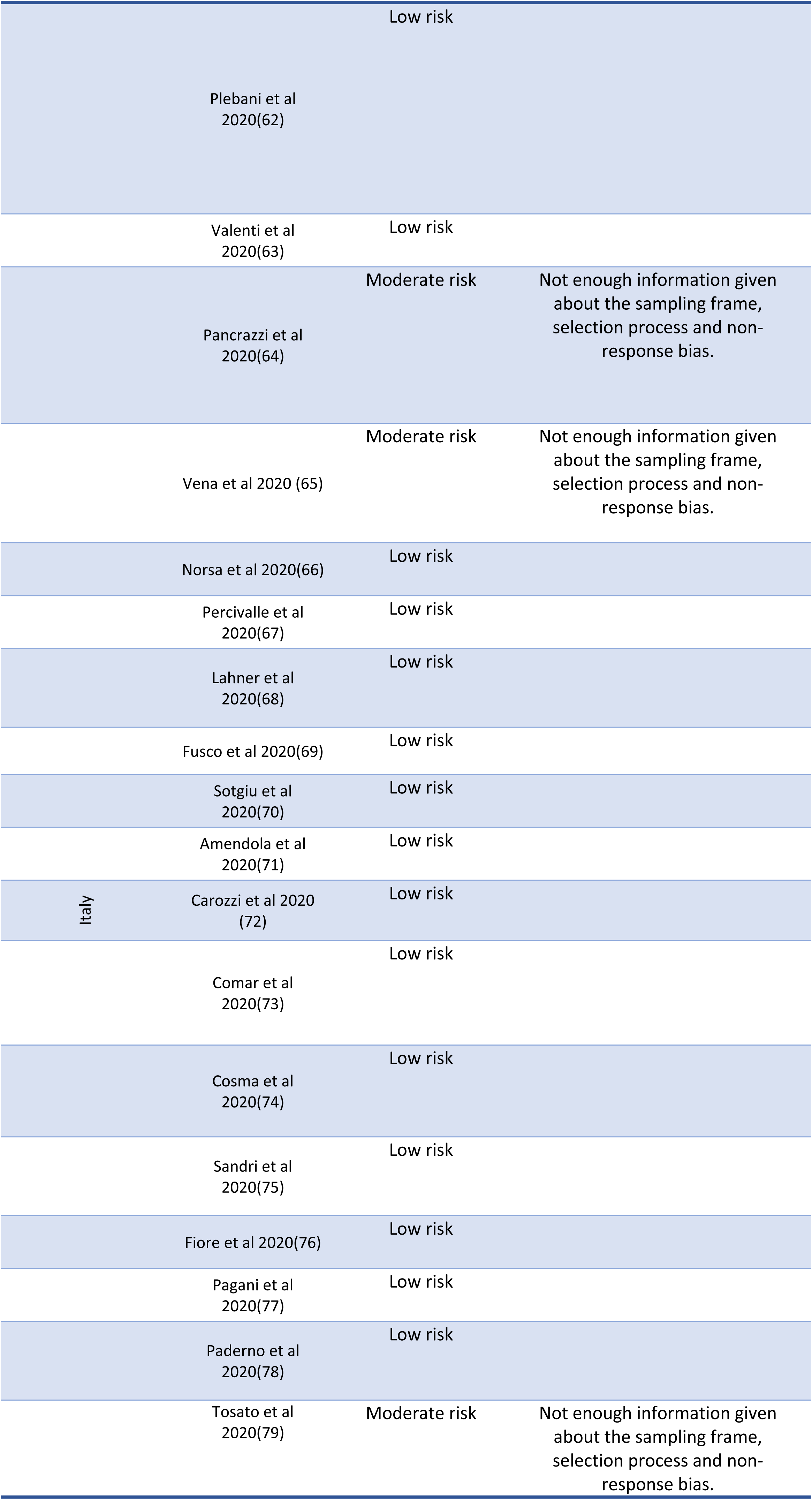
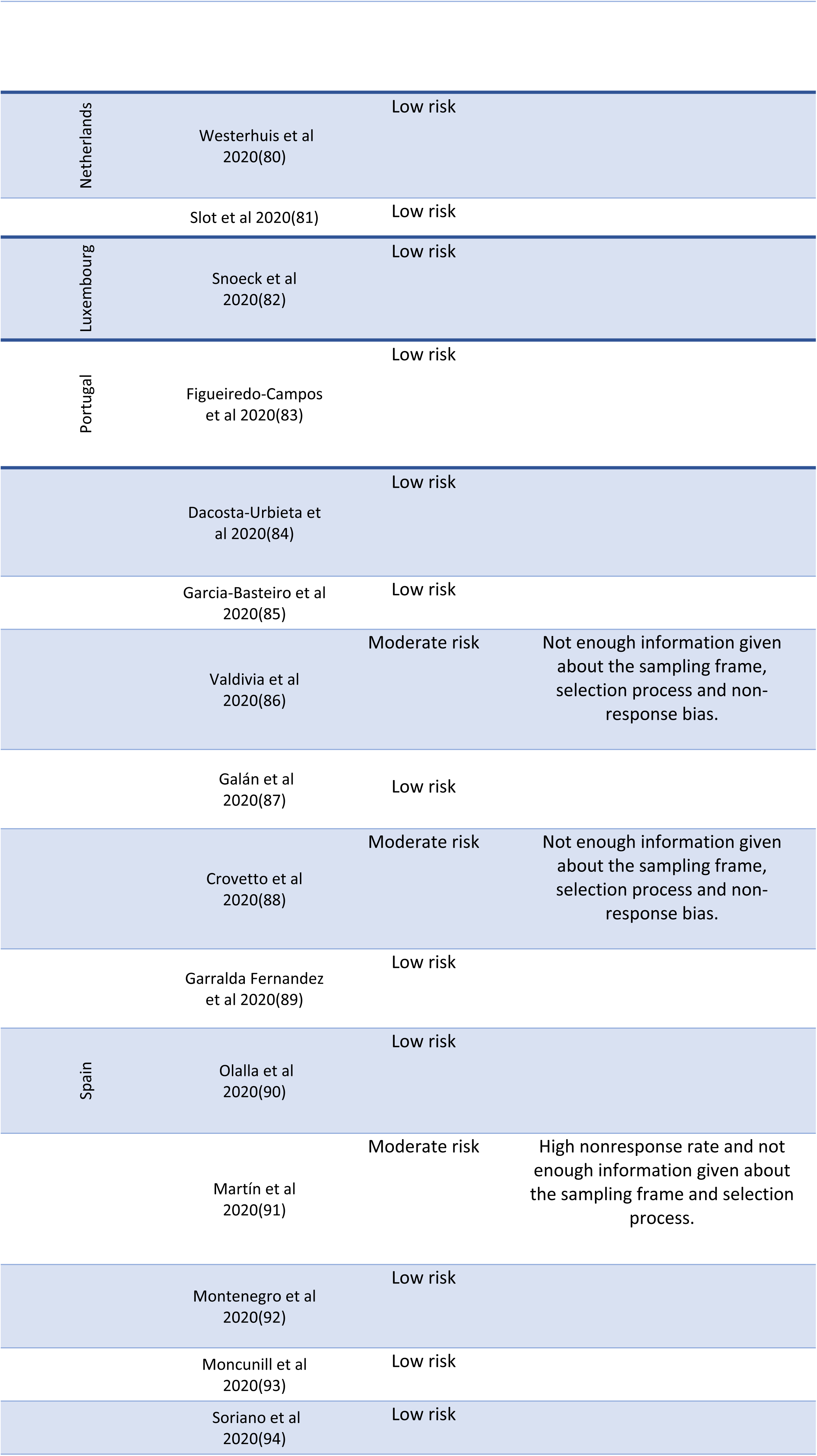
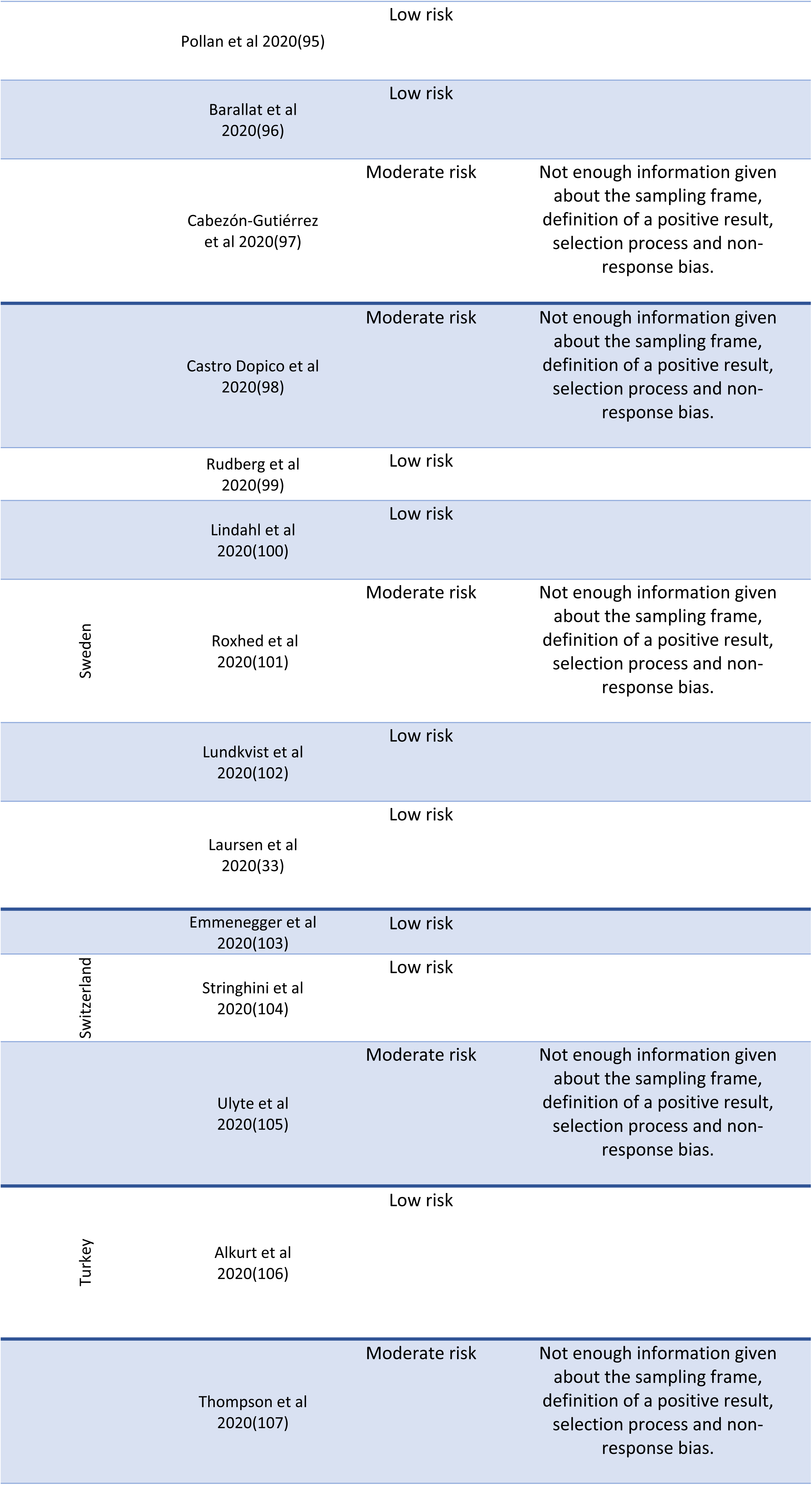
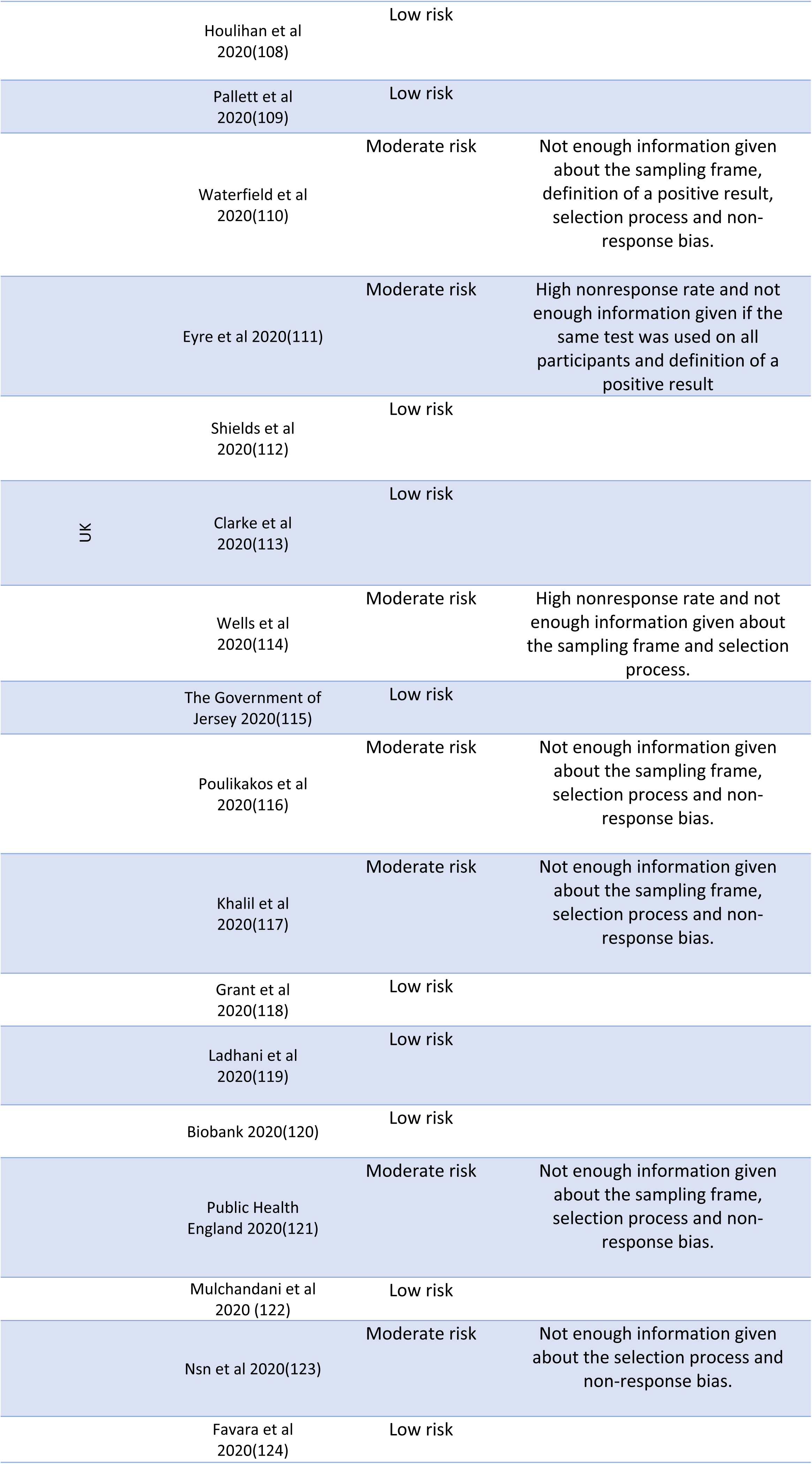
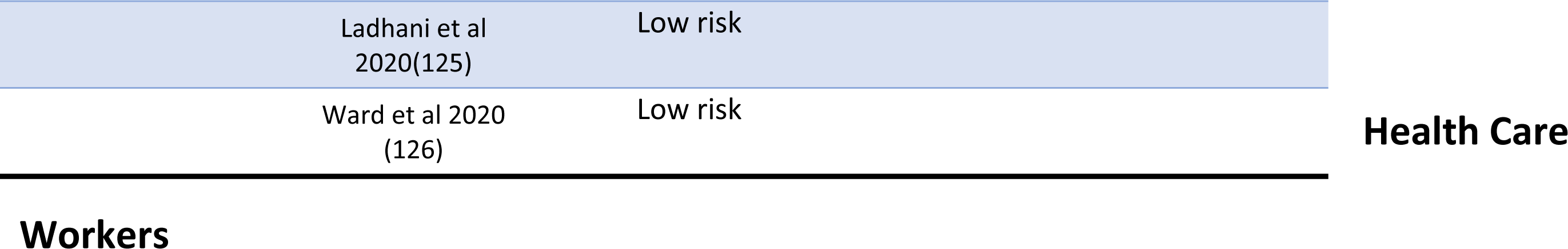
Assessment of the methodological quality of included studies

53 studies included seroprevalence data among HCWs. These studies included data from 13 countries in Europe and were conducted between February 2020 and August 2020 (Table 3).

**Table 3:**
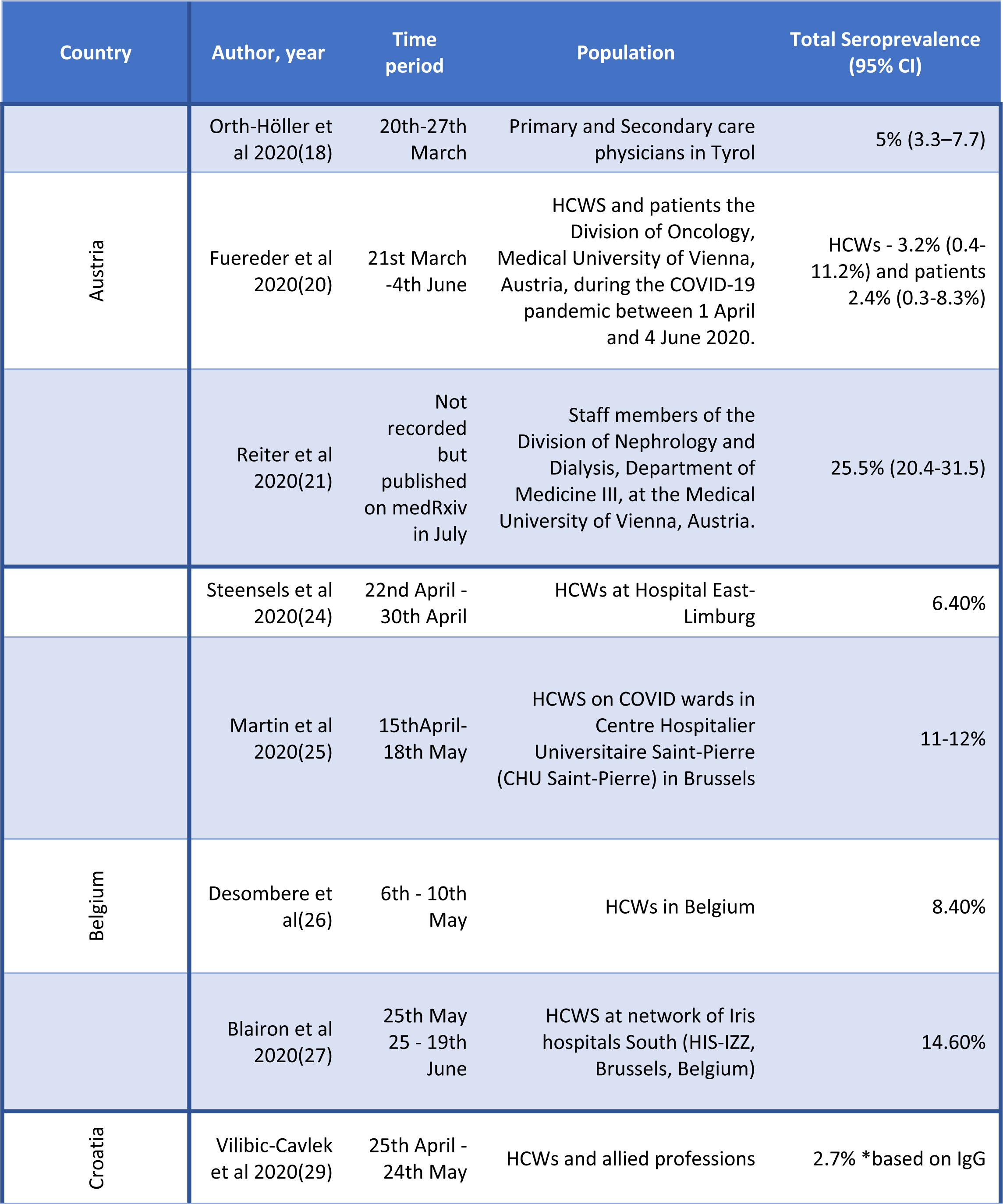
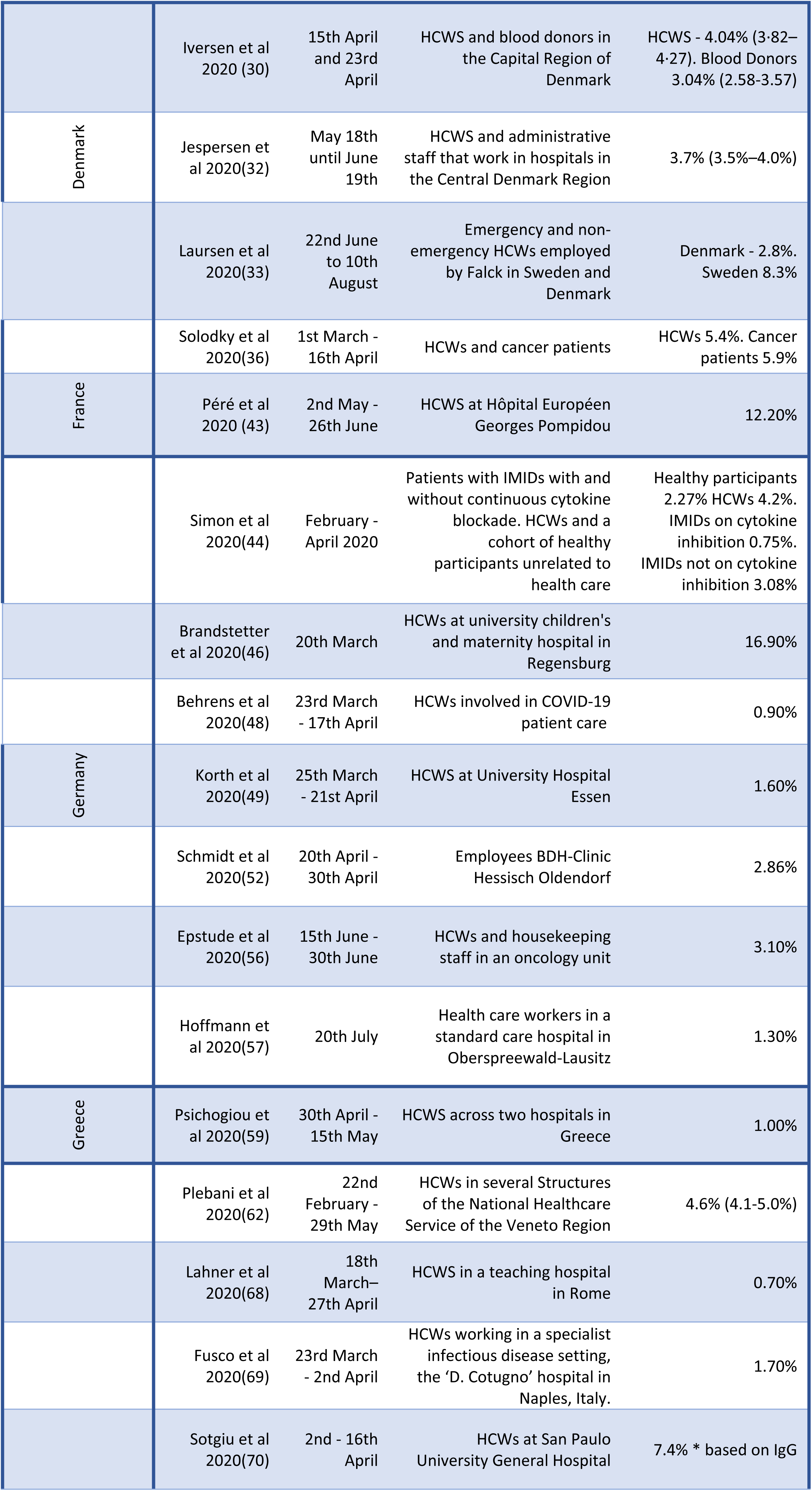
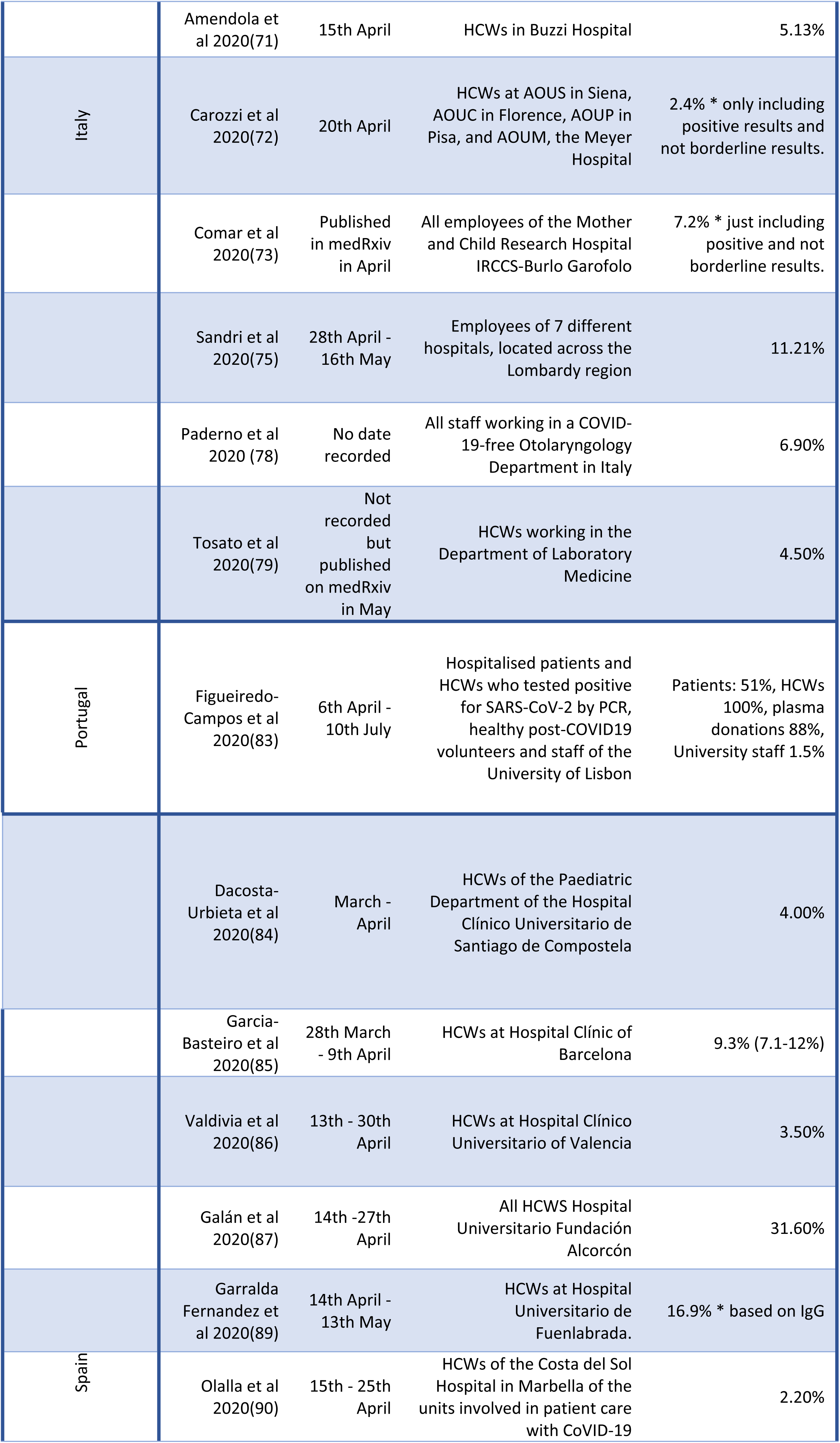
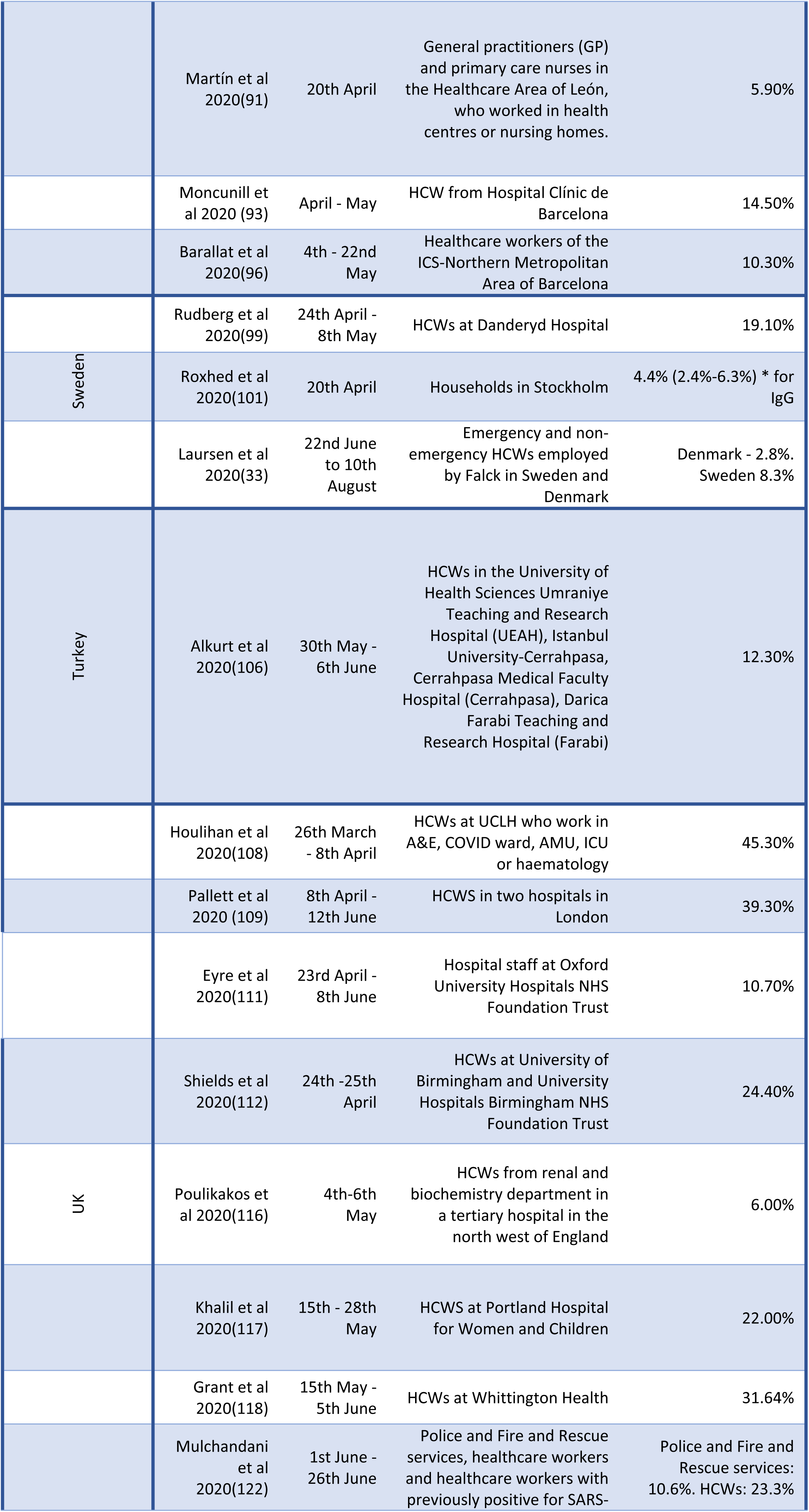
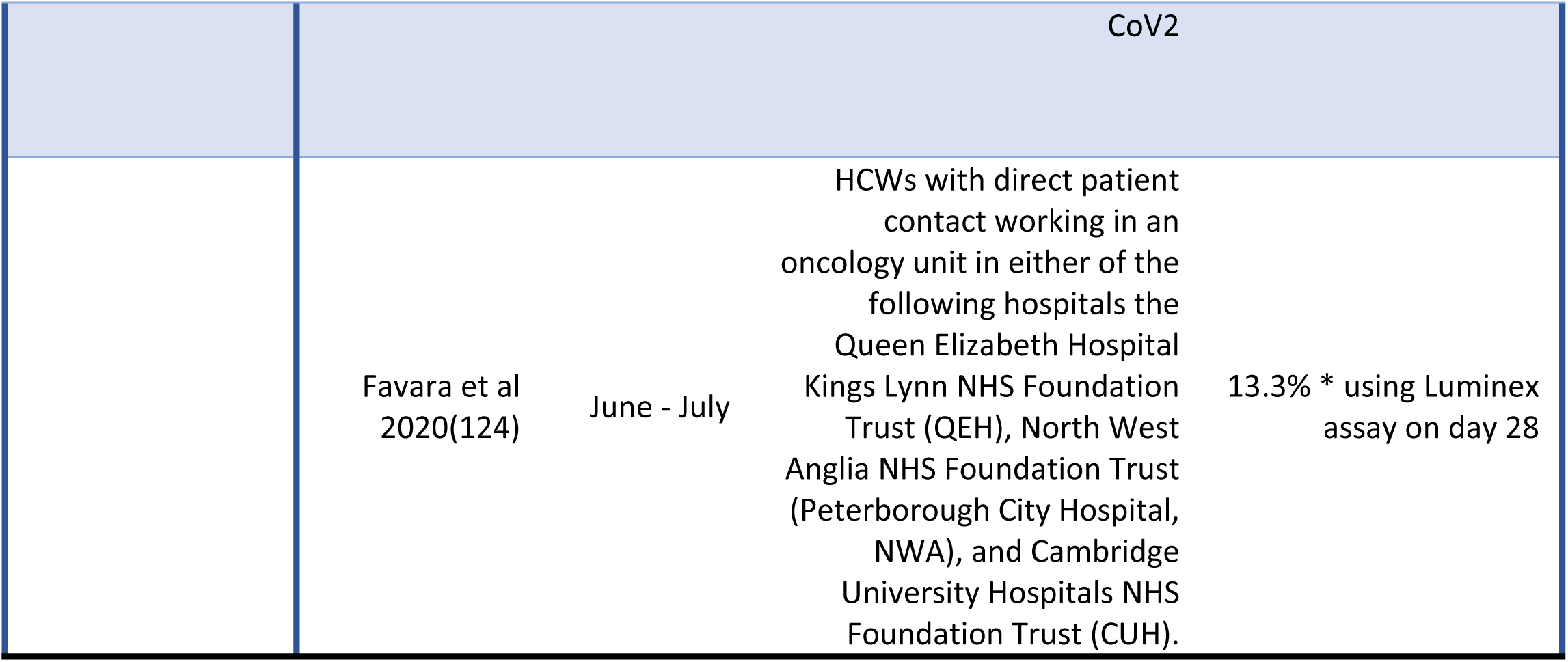
Studies reporting the seroprevalence among HCWs, dates the studies were conducted, population studied and the overall seroprevalence

The lowest seroprevalence was seen in a teaching hospital in Rome, Italy during the months of March – April 2020, reporting a seroprevalence of 0.7% based on IgG antibodies and 0% based on IgM (68). The highest seroprevalence (45.3%) was reported in March – April 2020 in a University Hospital in London (108). Figure 2, chart A shows the seroprevalence of HCWs by country overtime. The majority of studies report a seroprevalence < 10% between March – August 2020. A few studies predominately based in the UK report a seroprevalence 20-45% among HCWs during this time period (21,87,108,109,112,117,118,122).

**Figure 2:**
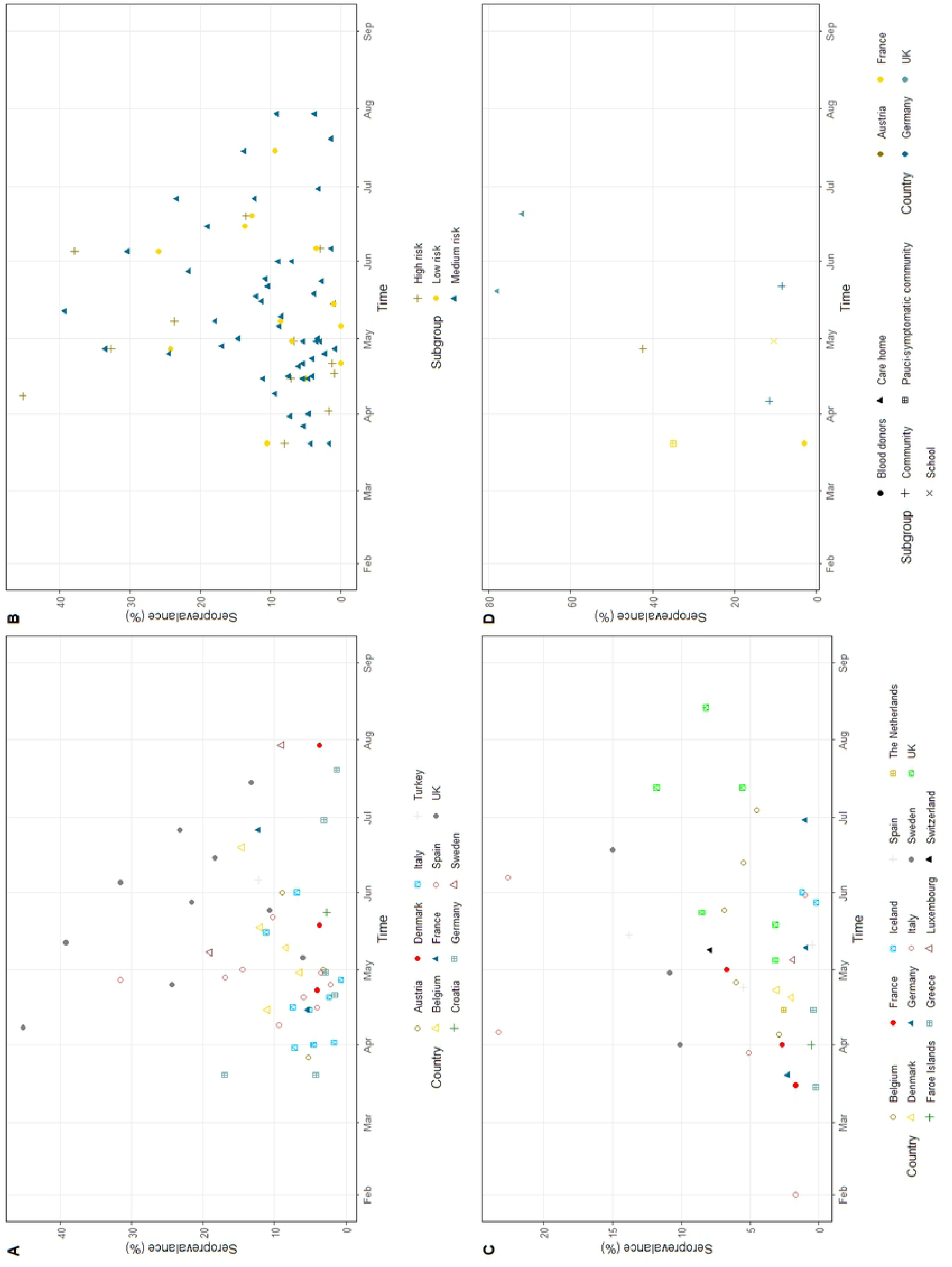
Chart A; shows the seroprevalence of HCWs by country overtime. Chart B: shows the seroprevalence of HCWs overtime stratified by risk group. Chart C: shows the seroprevalence of community studies overtime by country. Chart D: shows the seroprevalence of outbreak studies overtime stratified by country and subgroup.

Figure 2, chart B shows the seroprevalence of HCWs categorised by their risk of exposure to SARS-CoV-2 patients over time. All subgroups with a seroprevalence of >30% belonged to either medium or high risk. The majority (63/75) of the subgroups had a seroprevalence of less than 20% regardless of their risk.

There was no significant difference in seroprevalence amongst HCWs when stratified by country (Figure 3). There is a large amount of heterogeneity between the studies (I^2^ value = 99.3%, p = 0.00). There was no reduction in heterogeneity when moderate risk of bias studies were removed. Similarly, when the seroprevalence amongst HCWs was stratified by their risk of exposure to SARS-Cov-2 patients there was no significant difference (supplementary figure 1).

**Figure 3:**
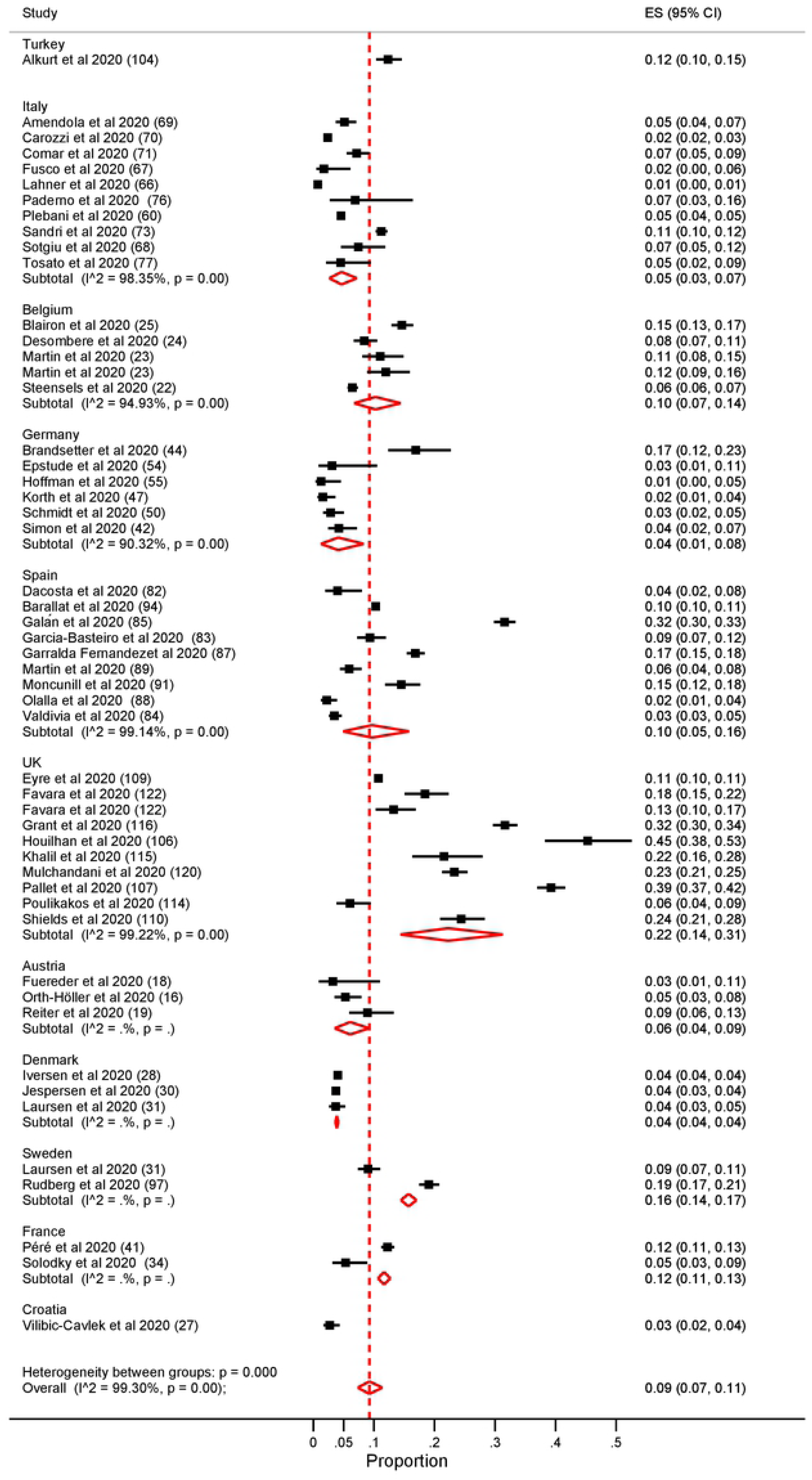
Forest plot of the seroprevalence among HCWs stratified by country.

## Community Studies

In total 33 studies were set in the community, spanning 13 countries. The studies collected data between February 2020 - August 2020 (Table 4). Ten of these studies collected samples from blood donors; four studies used residual serum samples from clinics, laboratories and hospital facilities; one study used tissues samples and the remaining studies were randomised population-based studies.

**Table 4:** Studies reporting the seroprevalence in a community setting, dates the studies were conducted, population studied and the overall seroprevalence

| Country | Author, year | Time period | Population | Total Seroprevalence (95% CI) |
| --- | --- | --- | --- | --- |
| Belgium | Herzog et al 2020(22) | 30 March -5 July | Residual sera from ten private diagnostic laboratories in Belgium. | 30 March – 5 April 2.90% (2.3-3.4%). 20 – 26 April 6% (5.1-7.1%). 18 – 25 May 6.9% (5.9-8). 8 – 13 June 5.5% (4.7-6.5%). 29 June – 4 July 4.5% (3.7-5.4%) |
|  | Iversen et al 2020(30) | 15th April and 23rd April | HCWS and blood donors in the Capital Region of Denmark | HCWS - 4.04% (3.82–4.27). Blood Donors 3.04% (2.58–3.57) |
| Denmark | Erikstrup et al 2020(31) | 6th April to 3rd May | Blood Donors in Denmark | 2% (1.8–2.2) |
|  | Petersen et al 2020(34) | 27th April – 1st May | Residents of the Faroe Islands | 0.6% (0.2%–1.2%) |
|  | Germain et al 2020(35) | 1st November 2019 to 16th March 2020 | All tissue donors at Lille Tissue bank | 1.70% |
|  | Grzelak et al 2020(37) | 20th March | A cohort of pauci-symptomatic individuals in Crepy-en-Valois. Blood donors from the Etablissement Français du Sang (EFS) in Lille (France) | Blood donors mean of 3%. Pauci symptomatic cohort mena of 32%. |
| France | Carrat et al 2020(38) | 1st April - 27th May | Residents from Ile-de-France (IDF) or Grand Est (GE) or Nouvelle-Aquitaine (NA) | 6.70% |
|  | Gallian et al 2020(39) | Last week of March to the first week of April | Blood Donors | 2.70% |
|  | Simon et al 2020(44) | February - April 2020 | Patients with IMIDs with and without continuous cytokine blockade. HCWs and a cohort of healthy participants unrelated to health care | Healthy participants 2.27% HCWs 4.2%. IMIDs on cytokine inhibition 0.75%. IMIDs not on cytokine inhibition 3.08% |
| Germany | Fischer et al 2020(45) | 9th March - 3rd June | Blood donors in three German federal states North Rhine-Westphalia, Lower Saxony and Hesse | 0.91% (0.58–1.24%) |
|  | Aziz et al 2020(53) | 24th April -30th June | Individuals enrolled in the Rhineland Study an ongoing community- based prospective cohort study in people aged 30 years and above in the city of Bonn, Germany. Group I all living participants who had been enrolled in the Rhineland Study until March 18, 2020.Group II individuals who were eligible for but had not yet participated in the Rhineland Study. | Group I: 0.97% (0.72–1.30). Group II: 1.94% (0.84–4.42) |
| Greece | Bogogiannidou et al 2020(58) | March and April | Individuals who visited the laboratories for a check-up, chronic disease follow-up or other reasons unrelated to COVID-19 in the whole of Greece | March: 0.24% (0.03–0.45%). April 0.42% (0.23–0.61%) |
| Iceland | Gudbjartsson et al 2020(61) | January - July | Two groups of qPCR-positive Icelanders and six groups of Icelanders who had not been qPCR-tested or who had been tested and had received a negative result | 4.22% |
|  | Valenti et al 2020(63) | 24th February - 8th April | Blood donors in Milan | 5.07% |
|  | Vena et al 2020(65) | March and April | Residents living in Liguria and Lombardi regions | 11.00% |
| Italy | Percivalle et al 2020(67) | 18th March - 6th April | Registered blood donors in Lodi Italy | 23.00% |
|  | Fiore et al 2020(76) | 1st May - 31st May | Blood donors in Apulia region, South Eastern Italy | 0.99% |
|  | Pagani et al 2020(77) | 18th May - 7th June | Residents of Castiglione D'Adda | 22.6% (17.2-29.1%) |
| The Netherlands | Slot et al 2020(81) | 1st - 15th April | Plasma donations | 2.60% |
| Luxembourg | Snoeck et al 2020(82) | 15th April - 5th May | Population of Luxembourg | 1.97% |
|  | Montenegro et al 2020(92) | 21 April to 24 April 2020 (Study population A) and from 29 April to 5 May 2020 (Study population B) | Population A: individuals registered in a primary health care centre, from a community area of Barcelona, Spain. Population B: Patients from GPs in Barcelona presenting with mild-moderate symptoms of COVID but no diagnosis of COVID | Population A: 5.47% (3.44–8.58%). Population B: 38.49% (34.78%–42.33%) |
| Spain | Soriano et al 2020(94) | April - May | Asymptomatic adults in Madrid | 13.80% |
|  | Pollan et al 2020(95) | 27th April - 11th May | Spanish households | 5% (4.7-5.4%) * based on POC testing |
|  | Castro Dopico et al 2020(98) | 30th March - 23rd August | Blood Donors and pregnant women in Stockholm | 13.7% (9.5-19.3%) * estimated by ENS Learner |
| Sweden | Roxhed et al 2020(101) | 20th April | Households in Stockholm | 4.4% (2.4%-6.3%) * for IgG |
|  | Lundkvist et al 2020(102) | 17th-18th June | Residents of Norra Djurgårdsstaden and Tensta in Stockholm | 15.00% |
| Switzerland | Stringhini et al 2020(104) | 6th April - 9th May | Residents from the Canton of Geneva | 7.90% |
|  | Thompson et al 2020(107) | 17th March - 19th May 2020 | Blood donors in Scotland | 3.17% |
|  | Wells et al 2020(114) | 27th April - 2nd June | Participants of the Twins UK Cohort Study | 12.00% |
|  | The Government of Jersey 2020(115) | 29th April - 5th May | Residents in Jersey over 16 years of age | 3.1% (+/- 1.3%) |
| UK | Biobank 2020(120) | 27th May - 14th August | UK Biobank participants | 8.2% (7.9%-8.7%) |
|  | Public Health England 2020(121) | May | Healthy adult blood donors, supplied by the NHS Blood and Transplant (NHS BT) | 8.5% (6.9-10%) * this is adjusted |
|  | Ward et al 2020(126) | 20th June - 13th July | Residents in England over the age of 18 years | 6% (5.8-6.1%) * adjusted for test performance and re-weighted for sampling |

The overall lowest seroprevalence was reported in Greece in March 2020 of 0.24% (0.03-0.45%) (58). The same study reported an increase in seroprevalence in April of 0.42% (0.23-0.61%). The overall highest seroprevalence was reported in Lodi, Italy during the months of April and March at 23% (67). The majority of studies reported an overall seroprevalence of less than 10% during the months of February – August 2020 (Figure 2C).

### Age

Many of the community studies report the seroprevalence among different age groups. There is significant heterogeneity between the results. In general, lower seroprevalences were reported at the extremes of age. Several studies report a higher seroprevalence among the over 50 age group (34,44,65,76,77,95). In contrast some studies report a higher seroprevalence in the less than 30 years age group, these include studies from Switzerland, The Netherlands, Denmark, France, Luxembourg and the UK (38,81,82,104,120,126).

### Gender

In the majority of community studies there was no significance difference identified by gender. However, two studies reported a significantly higher number of female participants having antibodies against SARS-CoV2 (38, 65). Carrat et al investigated the seroprevalence in three administrative regions of France Ile-de-France (IDF), Grand Est (GE) and Nouvelle-Aquitaine (NA) and reported a significant association of antibodies associated with the female gender only in Nouvelle-Aquitaine (38). Similarly, Vena et al report a significantly higher seroprevalence among female participants in five administrative regions in Italy (65).

### Blood donors

Many studies report the seroprevalence in blood donors as they are usually healthy individuals who represent the general population. There was a large variation in seroprevalence among blood donors between countries and overtime.

The lowest seroprevalence in blood donors was reported in Germany between March and June 2020 of 0.91% (0.58–1.24%) (45). In contrast Percivalle et al reported the highest seroprevalence amongst Italian blood donors in April, living in the Lodi Red Zone of 23.3% (67). The Lombardi Red Zone is an area of 10 municipalities that were put in total social and commercial lockdown from the 23^rd^ February 2020(67). The same study reports a seroprevalence of 1.67% in February 2020. In addition, a study in the South East of Italy reports a seroprevalence on 0.99% in May 2020(76).

Similar variations of estimates of seroprevalence were reported in blood donors in the UK. A study conducted in Scotland reported a seroprevalence of 3.17% between the months of March – May 2020(107). A seroprevalence of 8.5% (6.9-10%) was reported in blood donors across England in May (121).

## Children/ Schools

Six studies investigated the seroprevalence in school/university settings or among children only, across 5 different countries (41,55,60,105,110,125). Four of the studies examined the seroprevalence in schools (41,55,105,125). The lowest seroprevalence was reported in Germany; 0.6% among students in grade 8–11 and their teachers in 13 secondary schools in eastern Saxony between the months of May – June 2020 (55). The highest seroprevalence of 11.7% was reported in students and teachers across schools in England between June-July 2020 (10.5-13.3%) (125).

Fontanet et al investigated the seroprevalence among pupils and teachers in primary schools exposed to a SAR-CoV2 outbreak in Paris and reported an overall seroprevalence of 10.4% (41). They noted that 41.4% of infected children had asymptomatic infection compared to 9.9% of seropositive adults (41).

One study examined at the seroprevalence among university students and staff in Greece (60). They reported an overall seroprevalence of 1%, with no significant difference by age, gender, school or position (60).

## Outbreaks

Seven studies across four countries that investigated the seroprevalence following an outbreak of SAR-CoV2 (19,37,41,50,54,119,123) (Supplementary Table 1). Two of the studies were conducted in the UK and reported a high prevalence of antibodies against SARS-CoV2 in residents and staff in care homes/nursing homes where there had been a recent SARS-CoV2 outbreak (119, 123). They report a high prevalence of antibodies against SARS-CoV2. Ladhani et al estimated a seroprevalence of 77.9% (73.6-81.7%) and Nsn et al report a seroprevalence of 71.8% (119, 123).

In contrast four studies investigated the seroprevalences in the residents or blood donors in communities where there has been an outbreak (19,37,50,54). They report much lower rates of seroprevalence compared to nursing home outbreaks. Grzelak et al investigated the seroprevalence of pauci-symptomatic individuals in Crepy-en-Valois France and blood donors in the surrounding region following an outbreak; they reported a seroprevalence of 3% in blood donors and 32% in the pauci-symptomatic individuals (37). Similarly studies in Germany following community outbreaks report low rates of seroprevalence among residents. Streeck et al reported a prevalence of SARS-COV-2 antibodies of 13.6% and Weis et al reported a seroprevalence of 8.4% (50, 54). Figure 2, chart D shows the seroprevalence of these outbreak studies overtime.

## Pregnancy

Three studies examined the seroprevalence of SARS-CoV-2 in pregnant women (42,74,88). Two of these studies are conducted in Italy between April – June 2020 reported a prevalence of SARS-CoV-2 antibodies of 10.1% and 14.3% in pregnant women in their first trimester screening or at delivery (74, 88). The third study estimated a seroprevalence of 8% among pregnant women admitted to the delivery room in France in May 2020 (42). Mattern et al found that the seroprevalence among pregnant women was similar to that of the general public (42).

## Assays

In total 45 different commercial assays and 22 in-house assays were used. The majority of studies used more than one assay. Of the commercial assays 11 were enzyme-linked immunosorbent assay (ELISA), six were chemiluminescent microparticle immunoassays, two were based on flow cytometry and 26 were point of care tests (POC). The most commonly used commercial assay was the SARS-CoV-2 (IgA/IgG) ELISA EUROIMMUN Medizinische Labordiagnostik, Lübeck, Germany (supplementary table 2).

## Discussion

Our systematic review demonstrates a large variation in the seroprevalence of SARS-CoV-2 antibodies throughout Europe in the first half of 2020.

HCWs in the UK had a much higher seroprevalence compared to HCWs in the rest of Europe during the months of March and August 2020. There are nine studies which took place in UK and six of them reported a seroprevalence of more 20% among HCWs (108,109,112,117,118,122). In contrast Italy reports a low seroprevalence among HCWs. Of 10 studies among HCWs in Italy, nine reported a seroprevalence of SARS-CoV-2 antibodies of less than 10% (62,68–73,78,79). Both countries included studies from a mixture of high, medium and low risk HCWs and during this time both countries had high numbers of SARS-CoV-2 infections.

In health care settings the risk to HCWs of SARS CoV2 exposure will be determined by the COVID19 caseload coming though the facility and the application of infection control measures. Infection control practices in relation to personal protective equipment (PPE) may in part explain some of the differences.

Between European countries there are differences in the recommended PPE. The UK government guidelines on PPE include the use of eye/face protection, filtering facepiece class 3 (FFP3) respirator, disposable fluid-repellent coverall, and disposable gloves for aerosol-generating procedures and higher-risk acute care areas. For all inpatient ward settings eye/face protection, fluid-resistant (type IIR) surgical mask (FRSM), disposable plastic apron, and disposable gloves are recommended (130).

In comparison the National Institute of Health in Italy recommended that all HCWs wear a full-length gown with long sleeves, hairnet, goggles, gloves and surgical mask in the case of low-risk patients, and hairnet, googles or face-shield, FFP3 mask, water-resistant gown with long sleeves, and two pairs of gloves (second one covering the wrist of gown sleeves) for high-risk patients and SARS-CoV-2 positive patients (68).

Although the availability and differences in PPE across European countries may partly explain the difference in seroprevalence seen in HCWs, there are other factors that require consideration. For example, differences in public health strategies and the time of their implementation such as the public wearing face masks, closure of educational settings and other public facilities. Furthermore, differences in adherence to infection control measures such as hand hygiene and social distancing could also explain the difference in seroprevalence seen among HCWs across Europe.

Our systematic review found that in the majority of studies in Europe there was no difference in seroprevalence between female and male participants. Our findings are in keeping with a meta-analysis which showed there was no difference in the proportion of males and females with confirmed COVID-19(131).

However, there is strong evidence that males are more likely to have more severe disease compared to females. A meta-analysis looking at 92 studies world-wide concluded that male patients have almost three times the odds of requiring intensive treatment unit (ITU) admission and have higher odds of death compared to females (131). Similarly, Castro-Dopico et al reported that severe disease was associated with virus-specific IgA and that IgA responses were lower in females compared to males (98).

Throughout the current pandemic there has been debate on the role of children in the transmission of SARS-CoV-2 and the need for school closure to slow the pandemic. In this review three studies were in schools not involved in an outbreak of SARS-CoV-2. A study in Germany reported the lowest seroprevalence among students and teachers in a school of 0.6% considered by the authors to be in keeping with local surveillance data of the surrounding community (55). Ulyte et al reported that seroprevalence is inversely related to age in their school study (105). They conclude this could be due to the lack of social distancing among young children and differences in immune response (105). In contrast Ladhani et al reported no significant difference between the seroprevalence in students compared to staff (125). However, all studies concluded that there was no major transmission in schools and that the majority of children were asymptomatic or had mild symptoms (55,105,125). This review highlights the need for more school-based studies investigating the seroprevalence among staff and students to fully understand transmission dynamics and immune responses throughout the pandemic.

In studies conducted during local outbreaks there was a noticeable difference between those conducted in care/nursing homes compared to community and school settings. Those that took place in care/ nursing homes reported a seroprevalence as high as 77.9%, whereas those in a community setting reported a seroprevalence ranging from 3% - 42.4% and those in a school reported a seroprevalence of 10.4%(19,37,41,50,54,119) This large discrepancy could be attributable to the close proximity of care/nursing home residents, shared living spaces and the intimate care and handling of residents by staff.

## Limitations

This meta-analysis had several limitations. Firstly, of the 109 studies included in this review, not all of them could be included in sub-analysis as complete data sets could not be retrieved from every study and data quality was heterogeneous. In addition, the majority of studies were performed either in the UK, Italy, Spain or Germany. There was a large gap in studies being performed in Eastern European countries. Those studies performed in the UK predominately took place in the South of England. Furthermore, many of the studies were pre-print articles that had not undergone peer-review.

## Conclusion

This systematic review and metanalysis highlights substantial heterogeneity between countries, within countries, among professions, and among settings. This heterogeneity, in addition to indicating the general trajectory of the pandemic in different regions, will also be driven by a variety of other factors including governmental policies and restrictions, local guidelines and restrictions, availability of PPE, the time period when the study was conducted and serological test performance. Nevertheless, seroprevalence studies yield large amounts of useful, locally relevant information and should be regularly repeated as the pandemic evolves and local guidelines and restrictions change. As testing standardizes and new studies are reported they will also help identify different national experiences across Europe and provide a means to distil best pandemic control practices for the future.

Finally, as new variants of SARS-CoV-2 now emerge, and many countries prepare for future waves it is vital that regular seroprevalence studies are conducted to aid control in the context of vaccine implementation.

## Author Contributions

**NMV –** wrote the protocol, develop search terms, conducted searches, screened studies, extract data, conducted data analysis and meta-analysis and wrote the final draft.

**DH –** wrote the protocol, develop search terms, conducted data analysis and meta-analysis and proofread the final draft.

**BS-** wrote the protocol, develop search terms, conducted searches, screened studies and proofread the final draft.

**AK –** extracted data and proofread the final draft.

**CN –** helped with writing and proofreading the final draft

**NF –** wrote the protocol, develop search terms, helped to conduct data analysis and meta-analysis and proofread the final draft.

All authors have approved and read the final manuscript.

## Acknowledgements

NC and DH are affiliated to the NIHR Health Protection Research Unit (HPRU) in Gastrointestinal Infections at University of Liverpool, in partnership with Public Health England (PHE), in collaboration with University of Warwick. NC and DH are based at The University of Liverpool. NF and DH are affiliated to the NIHR HPRU Emerging and Zoonotic Infections, a partnership between PHE, the University of Liverpool in collaboration with the Liverpool School of Tropical Medicine and the University of Oxford. The views expressed are those of the author(s) and not necessarily those of the NIHR, the Department of Health and Social Care, HPRU or PHE.

## Supplementary Files

*Supplementary Table 1: Studies reporting the seroprevalence in an outbreak setting, dates the studies were conducted, population studied and the overall seroprevalence*

*Supplementary Table 2: Commercial assays and their specificity and sensitivity according to the manufacturer. In some cases, the manufacture’s specificity and sensitivity were unable to be found, so evaluation study data was used instead.*

*Supplementary figure 1: Forest plot of the seroprevalence among HCWs stratified by risk of exposure to SARS-CoV-2 patients.*

